# Abiotic and biotic stress activate the unfolded protein response in grapevine, with IRE1/bZIP60 signaling contributing to *Botrytis cinerea* resistance

**DOI:** 10.1101/2025.08.11.669611

**Authors:** Tania Marzari, Emma Poilvert, Cécile Blanchard, Karine Palavioux, Agnès Klinguer, Hugo Hassangatebois, Alix Martinet, Aurélien Henry, Benoit Poinssot, Mathieu Gayral

## Abstract

A wide range of stresses can lead to the accumulation of unfolded or misfolded proteins in the lumen of the endoplasmic reticulum (ER), a condition known as ER stress. To restore proteostasis, eukaryotic cells activate a signaling network called the unfolded protein response (UPR). In *Vitis vinifera*, two arms of the UPR have been identified: the IRE1/bZIP60 arm and the bZIP17 arm. Notably, no putative ortholog of bZIP28 has been found in *V. vinifera*, whereas this is the case for *Arabidopsis thaliana*. We demonstrated that *VvbZIP60* mRNA undergoes unconventional splicing upon treatment with dithiothreitol (DTT) and tunicamycin (TM), both classical ER stress inducers. In addition, inhibition of IRE1 RNase activity directly decreases IRE1-bZIP60-related gene expression. We observed a strong transcriptional activation of UPR-related genes in response to heat stress which induced the re-localization of both VvbZIPs to the nucleus. Surprisingly, only VvbZIP17 responded to osmotic stresses. Grapevine is also subject to a broad range of microbial challenges, including pathogens such as *Plasmopara viticola* and *Botrytis cinerea*. Both pathogens triggered UPR genes activation in grapevine leaves. Interestingly, inhibition of IRE1 RNase activity induced higher susceptibility to *B. cinerea*. Collectively, these findings suggest that the IRE1/bZIP60 and bZIP17 arms of the UPR are not only structurally conserved in grapevine but are also transcriptionally responsive to a variety of abiotic and biotic stresses.

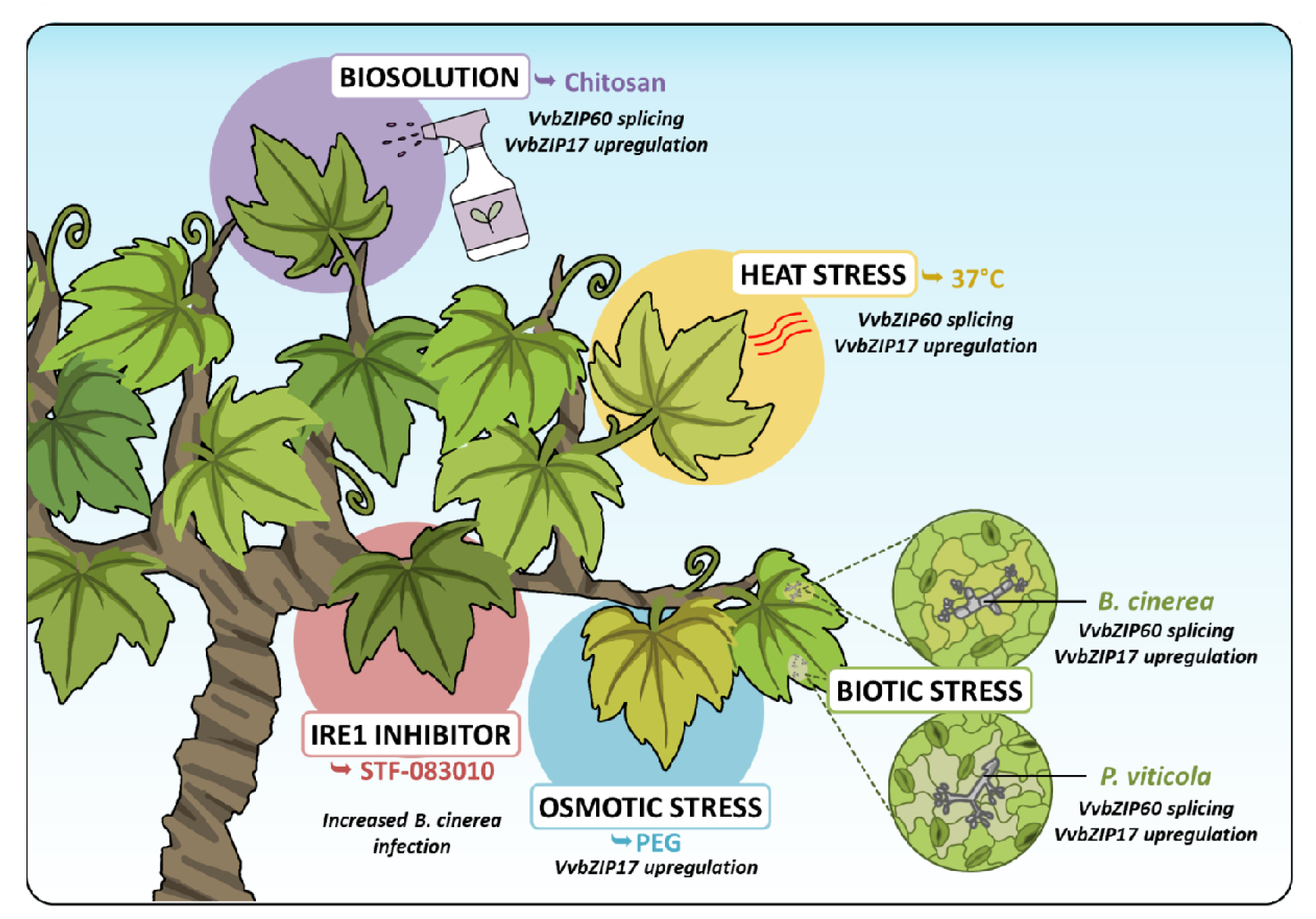

## 1. Introduction

Grapevine cultivation is one of the most significant agricultural activities worldwide, it covers approximately 7 million hectares of the total rural surface and holds immense economic weight on the international market generating over 33.8 billion euros in wine production, estimated to be around 227 million hectoliters (OIV, International Organization of Vine and Wine, 2025). However, this sector faces increasing challenges due to extreme climatic conditions and fungal diseases, which have significantly affected yield and fruit quality (Adrian et al., 2024). As a perennial crop, grapevine growth relies on stable climatic conditions, including optimal temperatures, adequate water supply and frost-free periods (Palliotti and Poni, 2015; Zapata et al., 2017). Unfortunately, maintaining these conditions is becoming increasingly difficult, thereby raising the challenges of grapevine cultivation.

Plants have evolved a range of tolerance mechanisms to cope with both abiotic and biotic stress conditions and the unfolded protein response (UPR) is one of them. Acting as a sentinel for adverse conditions, the UPR pathway allows the maintaining of cellular homeostasis controlling protein folding and limiting accumulation of misfolded and/or unfolded proteins in the endoplasmic reticulum (ER) (Howell, 2013). The adaptation to changing conditions or pathogen attacks triggers extensive transcriptional reprogramming, which can overwhelm the ER’s capacity to manage protein folding properly. Notably, environmental factors including heat, salt or drought (Liu et al., 2007a; Valente et al., 2009; Deng et al., 2011) and interactions with microorganisms (Moreno et al., 2012; Gayral et al., 2020; Qiang et al., 2021; Blanchard et al., 2025; Marzari et al., 2025a) can intensify the workload of the ER beyond its capacity. This leads to the accumulation of misfolded and unfolded proteins within the ER lumen, a condition referred as ER stress (Howell, 2013). In response to this accumulation, the ER activates the UPR pathway to balance its protein-folding and degradation capacities with the cell’s needs (Walter and Ron, 2011). When managed effectively, short-term ER stress can be alleviated, allowing the ER machinery to recover. However, if ER stress persists, it can lead to autophagy, programmed cell death (PCD) or even the death of the entire plant (Iwata and Koizumi, 2005a; Yang et al., 2014).

In plants, two arms of the UPR pathway have been described in the model plant *Arabidopsis thaliana,* the ER-transmembrane inositol-requiring enzyme 1 (IRE1) / basic leucin zipper 60 (bZIP60) arm and the membrane-associated transcription factors (TFs) bZIP17/bZIP28 arm (Liu et al., 2007a; Nagashima et al., 2011; Hollien, 2013; Howell, 2013). Interestingly, in the *A. thaliana* genome, three isoforms of IRE1 have been described: AtIRE1A, AtIRE1B and AtIRE1C (Deng et al., 2011; Koizumi et al., 2001). While AtIRE1C is primarily involved in development and gametogenesis (Mishiba et al., 2019; Pu et al., 2019), AtIRE1A and AtIRE1B are particularly active during ER stress to mediate the unconventional splicing of bZIP60 mRNA resulting in a 23-pb elimination and a 3’ frameshift, encoding an UPR-specific TF lacking its transmembrane domain and presenting an additional nuclear localization signal (NLS) (Nagashima et al., 2011; Zhang et al., 2015). The translated spliced form of *AtbZIP60* (bZIP60s) is then translocated to the nucleus, where it activates downstream UPR responsive genes (Deng et al., 2011). In *A. thaliana*, the unconventional splicing of *bZIP60* mRNA is incomplete, with the unspliced form (bZIP60us) consistently present in high amounts. Depending on stress conditions, the splicing reaches its peak between 0.5 and 2 hours, after which the level of bZIP60 decreases (Parra-Rojas et al., 2015). The IRE1 activation occurs through conformational changes triggered by lateral oligomerization in the ER membrane (Belyy et al., 2022). While several models explain IRE1 dimerization in yeast and mammals (Gardner and Walter, 2011; Dawes et al., 2024), these mechanisms remain unclear in plants. One model suggests that BiP (binding protein), a predominant ER-resident Hsp70 chaperone normally associated with IRE1, is released under ER stress allowing IRE1 self-dimerization thus initiating the bZIP60 splicing (Bertolotti et al., 2000; Kimata et al., 2003). Another model proposes that IRE1’s luminal domain directly interacts with unfolded proteins (Gardner and Walter, 2011). Nowadays, the components of the IRE1-bZIP60 pathway and the splicing process have been identified in the model plant Arabidopsis and few other crop species, notably after treatment with ER stress inducers such as tunicamycin (TM) and dithiothreitol (DTT) which inhibit glycosylation and disulfide bond formation, respectively (Deng et al., 2011; Nagashima et al., 2011; Hayashi et al., 2012). In addition to its splicing activity, IRE1 performs a second function called Regulated IRE1-Dependent Decay (RIDD), in which it degrades mRNAs localized to the ER membrane in association with ribosomes (Li and Howell, 2021). RIDD plays a pivotal role during pathogen infection (Blanchard et al., 2025) and is a key regulator of UPR-induced autophagy (Bao et al., 2018). Moreover, IRE1 acts on the ubiquitin–proteasome system through the plant-specific E3 ligase PIR1 to regulate ER stress-induced programmed cell death (PCD) (Ko et al., 2023).

The second arm of the UPR pathway is mediated by the ER membrane-associated TFs bZIP17 and bZIP28 in *A. thaliana*. Under normal conditions, the luminal domain of AtbZIP28 is associated with BiP chaperone proteins (Srivastava et al., 2013). Under stress conditions, AtbZIP28 dissociates from BiP, and along with AtbZIP17, is translocated to the Golgi apparatus, where it undergoes sequential cleavage by site-1 protease (S1P) and site-2 protease (S2P). Cleavage by S2P releases the N-terminal domain of bZIP17/bZIP28, allowing their relocation to the nucleus to activate stress-responsive gene expression (Liu et al., 2007a, 2007b). However, the exact mechanisms for the activation of this UPR arm remain under investigation. For example, studies suggest that AtS1P may not be essential for activating AtbZIP28 (Iwata and Koizumi, 2005a; Iwata et al., 2017) and there is no direct evidence of a bZIP17/BiP association/dissociation during ER stress. Despite this, both bZIP17 and bZIP28 have been shown to be active under different stresses such as heat, drought and salt stress in various plant species (Yang et al., 2013; Silva et al., 2015; Ramakrishna et al., 2018). However, functional differences between these two TFs have been observed in Arabidopsis. For instance, AtbZIP28 has been described as the more prominent activator of the canonical UPR induced by proteotoxic stress, whereas AtbZIP17, together with AtbZIP28, plays a larger role in sustaining the expression of growth-related genes and is more closely associated with environmental stress responses than with canonical ER stress signaling (Kim et al., 2018, 2022). In addition, in *A. thaliana*, *AtbZIP28* is strongly induced by the ER-stressor DTT and TM (Tajima et al., 2008) whereas AtbZIP17 is more closely associated with salt stress responses (Liu et al., 2007a; Cho, 2024) and development (Bao and Howell, 2017; Gao et al., 2022). Interestingly, similar distinctions have been observed in rice: the orthologs of AtbZIP28 and AtbZIP17, respectively OsbZIP39 and OsbZIP60 (Wakasa et al., 2011), respond in a same manner as their Arabidopsis counterparts. Specifically, the transcript of *OsbZIP39* is induced by DTT and TM, but not by NaCl, suggesting that it may be a functional ortholog of AtbZIP28 rather than AtbZIP17 (Takahashi et al., 2012). Nevertheless, the precise functional differentiation between bZIP17 and bZIP28 remains not completely understood, reflecting both the conserved and lineage-specific regulatory complexity of UPR signaling across plant species.

Upon ER stress, bZIP TFs bind to the promoters of UPR target genes, such as BiPs through ER stress-responsive cis-regulatory elements (CREs) (Iwata and Koizumi, 2005b; Liu et al., 2014; Ko and Brandizzi, 2022a; Geng et al., 2026). These CREs are conserved DNA motifs that serve as binding sites which mediate transcriptional upregulation of genes involved in protein folding, quality control, and ER homeostasis (Oh et al., 2003). For example, in Arabidopsis BiP promoters, well-known CREs have been identified, particularly the unfolded protein response element (UPRE-II and UPRE-III) and the ER stress-response element (ERSE-I and ERSE-II) (Liu et al., 2014; Ko and Brandizzi, 2022a, 2024). Beyond these CREs, the activation of UPR-targeted genes is fine-tuned by co-regulators of bZIP28 and/or bZIP60, including HY5 (Nawkar et al., 2017), GBF2 (Ko and Brandizzi, 2022b) and ABI5 (Ko et al., 2023).

Although the UPR is not yet fully understood, extensive studies in various plant species have provided valuable insights into its regulatory mechanisms. These studies have revealed differences in UPR activation and composition across species under various stress conditions (Manghwar and Li, 2022; Rana et al., 2026). However, most research have primarily focused on few abiotic stresses like heat and salt, and even less studies examined biotic factors. Among these, the first evidence linking the UPR to plant immunity was obtained in *A. thaliana* and tobacco plants challenged with bacterial pathogens (Moreno et al., 2012; Arraño-Salinas et al., 2018) and viruses (Adhikari et al., 2025). More recently studies were expanded also to fungal or oomycetes infection (Qiang et al., 2021; Blanchard et al., 2025; Marzari et al., 2025a). In addition, *A. thaliana* is the most widely used model, limiting our understanding of UPR’s role during pathogen attacks in different contexts.

In this study, we aimed to characterize the UPR in grapevine (*Vitis vinifera*), a crop of major economic importance worldwide. Through phylogenetic analysis, we identified putative members of the two main UPR arms and analyzed the transcriptional profiles of *VvIRE1, VvbZIP60, VvBiP3*, and *VvbZIP17* in response to abiotic and biotic stresses. We showed that IRE1 mediates the unconventional splicing of *VvbZIP60* mRNA and that heat-induced ER stress triggers the re- localization of both VvbZIPs to the nucleus. Finally, we showed that the well-known grapevine pathogens *Botrytis cinerea* and *Plasmopara viticola* induce the UPR, and that the IRE1-bZIP60 arm is involved in grapevine susceptibility to *B. cinerea*.

## 2. Materials and methods

### 2.1. Plant materials

*Vitis vinifera* cv Marselan cell culture and grapevine herbaceous cuttings were grown as previously described (Brulé et al., 2019; Roudaire et al., 2023; Marzari et al., 2025b).

### 2.2. Phylogenetic analysis

Protein sequences of bZIP17, bZIP28, bZIP60, BiP and IRE1 were retrieved using tblastx searches, in which Arabidopsis family members were used as query sequences against protein sequences from plant species detailed in Supplementary Table S1. In addition, orthologous gene families were identified with OrthoFinder (v2.5.4). The bZIPs sequences were confirmed in different databases such as EnsemblPlants (https://plants.ensembl.org/index.html), NCBI (https://www.ncbi.nlm.nih.gov/) and UNIProt (https://www.uniprot.org/). Multiple full-length protein sequences alignment (ClustalW method) and a maximum-likelihood phylogenetic tree with 1000 bootstraps were used choosing the Maximum Likelihood method and JTT model (Jones et al., 1992). Alignment and phylogenetic analysis were performed with MEGA X software (Tamura et al., 2021).

### 2.3. Cell suspension treatments

Grapevine cell suspension was subjected to different abiotic stresses at the following final concentration: 2 mM dithiothreitol (DTT), 2 μg/mL tunicamycin (TM), 1% polyethylene glycol 8000 (PEG), 50 mM sodium chloride (NaCl), 10 nM of XcFlg22 and 0.2 mg/mL of hexamer of chitin (CO6, GLU436), chitosan (CHITO, GLU426) and tetramer of chitin (CO4; GLU434). For the IRE1 inhibition assay, STF-083010 (Sigma) was used at 60 μM. All treatments were diluted in water with the exception of the tunicamycin and STF-083010 which were dissolved in dimethyl sulphoxide (DMSO). For heat and cold stress, 20 mL of grapevine cell suspension contained in a 50 mL flask were put in a water bath at 37°C or put on a shaker plate in a cold room at 4°C. In parallel, control cell suspension was also treated with the corresponding volume of water or DMSO and collected at the same time point as the treated samples.

### 2.4. Pathogen infection

For *Botrytis cinerea* infection assays and STF-083010 inhibition assay, leaf disks were inoculated as previously described (Marzari et al., 2025b). For *Plasmopara viticola* infection assays, the lower leaf surface of grapevine greenhouse plants were sprayed as described in Marzari et al. 2025b.

### 2.5. VvbZIP60 splicing assay

For gel-based bZIP60 splicing assay, 5 ng of ADNc were amplified by PCR with specific primers flanking the unconventional intron, VvbZIP60_F and VvbZIP60_R (Supplementary Table S2). PCR conditions for amplification were: initial denaturation: 3 min at 95°C; 45 s at 95°C, 30 s at 63°C and 30 s at 72°C during 35 cycles; final extension 5 min at 72°C. Subsequently, PCR products were resolved by gel electrophoresis on agarose (3% w/v) (Invitrogen) using TAE 0.5X as running buffer.

### 2.6. Real-time quantitative reverse transcription polymerase chain reaction (qRT-PCR)

Grapevine cell suspensions were treated and harvested 1 and 3 h after elicitors and abiotic stresses. For IRE1 inhibition assay, cell suspension was treated with STF-083010 either together with the elicitor or 10 or 60 min before DTT or TM treatment. For pathogens infection test, leaf samples were harvested 5- and 7-days post *P. viticola* infection and after 24 – 48 and 72 hours post *B. cinerea* infection. Total plant RNAs were extracted using the SV Total RNA Isolation System kit (Promega) with DNAse treatment according to the supplier instructions for grapevine cell suspension and with the Plant/Fungi Total RNA Purification Kit (NORGEN) with some modifications for leaf disks. Reverse transcription (RT) was performed as described previously (Marzari et al., 2025b). Expression data were analyzed from the average of threshold cycle (CT) values taking into consideration the efficiency (E) of each reaction calculated by the LinRegPCR quantitative PCR data analysis program (Ruijter et al., 2009) and normalized on two references genes. Relative gene expression was assessed according to the Common Base Method (delta delta Cq - Ganger et al., 2017). The sequences of primers used can be found in Supplementary Table S2.

### 2.7. **Overexpression vectors for subcellular localization of UPR proteins**

Overexpression vectors for subcellular localization of UPR proteins were generated using the GoldenBraid 2.0 cloning system. Coding sequences and individual modules were either PCR-amplified and domesticated into the pUPD2 vector, or directly synthesized and cloned into a pDGB3α1 vector. The ER was visualized using the 35S:Kar2-CFP-HDEL:tNOS construct. VvbZIP60 and VvbZIP17 were N-terminally fused to RFP and expressed under the control of the 35S promoter and tNOS terminator. Final constructs used for subcellular localization assays were pDGB3Ω1_35S:mRFP1-VvbZIP60:tNOS_35S:Kar2-CFP-HDEL:tNOS and pDGB3Ω1_35S:mRFP1-VvbZIP17:tNOS_35S:Kar2-CFP-HDEL:tNOS.

### 2.8. Microscopy

CFP and mRFP1 signals were acquired sequentially 48 h post-transfection of *N. benthamiana* leaves, using a Leica SP8 confocal microscope fitted with an HC PL APO CS2 63x/1.40 oil-immersion objective. CFP was excited at 448 nm and emission collected between 460–500 nm; mRFP1 was excited at 552 nm and emission collected between 570–630 nm. Images were processed with LAS X (Leica), Fiji, and Inkscape software.

### 2.9. Cell viability

Grapevine cell suspension was stained with Fluorescein Diacetate (FDA) probe diluted at 20 μg/mL in 0.26% acetone then observed under an epifluorescence microscope (λexc: 450-490 nm, λem: 515 nm (LP), magnification x20) (Leitz, model DMRB). For each treatment condition, at least 200 cells were counted twice from three independent biological replicates.

### 2.10. *B. cinerea* growth assay

The effect of STF-083010 on *Botrytis cinerea* growth was assessed using a Bioscreen C MBR instrument (Oy Growth Curves Ab Ltd, Finland). Conidia were harvested from sporulating cultures and adjusted to a final concentration of 2×10^5^ spores/mL in ¼ PDB medium. Four conditions were tested: *B. cinerea* alone, *B. cinerea* supplemented with STF-083010 at 60 µM, STF-083010 alone and DMSO control. For each condition, 300 µL of the conidial suspension (or medium alone, for the negative control) was dispensed into the wells of a honeycomb 100-well microplate, with 3 technical replicates per condition. Plates were incubated in the Bioscreen C at 28 °C and absorbance was measured at 492 nm every 2 h over a total period of 62 h. Fungal growth was expressed as absorbance values over time, and growth curves were compared between conditions.

### 2.11. *P. viticola* zoospore encystment

Direct effects on zoospore encystment were assessed by microscopic visualization using aniline blue. Grapevine disks were treated with STF-083010 at 60 µM on the abaxial sides by flotation for 1 h. Disks were then inoculated on the abaxial side with a *P. viticola* sporangia suspension in water (10 spores/mL) and placed in a humid crystal box. At 24 h post-inoculation, leaf discs (8 mm diameter, at least 10 per condition) were collected and fixed in absolute methanol overnight to remove chlorophyll pigments. Discs were cleared in chloral hydrate (1 mg/mL) until fully transparent, rinsed once briefly and twice for 10 min in 0.1 M phosphate buffer (pH 8-9.3), and stained overnight in a freshly prepared aniline blue solution. Discs were mounted between slide and coverslip with a drop of aniline blue solution and observed under an epifluorescence microscope (10× objective; excitation 340-380 nm, emission 425 nm, LP filter). Three representative photographs were taken per disc, for a total of 10 discs per condition per biological replicate, using NIS-Elements BR 2.30 software. Zoospores were counted manually to quantify the proportion of encysted zoospores.

### 2.12. Statistical analysis

For qRT-PCR analysis, ΔCq values from treated or infected samples were compared with those of the control condition at the same time point using a one-way ANOVA followed by Tukey’s HSD test if normality was assumed otherwise a Kruskal-Wallis test followed by a Dunn post doc test was used. Data are presented as log2 fold change relative to the control condition. For sporulation and encystment assay, a linear mixed model (LMM; Gaussian distribution) was used, with treatment as a fixed effect and biological replicate as a random intercept. For necrosis area, a generalized linear mixed model (GLMM) with a Gamma distribution and a log link function was used, with treatment as a fixed effect and biological replicate as a random effect.

## 3. Results

### 3.1. Grapevine conserves key components of both UPR arms

A previous study identified 55 members of the bZIP family in the grapevine genome, including VvbZIP24 (here renamed VvbZIP17) and VvbZIP41 (here renamed VvbZIP60) (Liu et al., 2014). To characterize potential TFs and other actors involved in ER stress in *Vitis vinifera*, we conducted a phylogenetic analysis using reference sequences from *A. thaliana* and their relative orthologs in mammals and yeast. Concerning the bZIP TFs, we retrieved 28 protein sequences for bZIP60 and 50 protein sequences for bZIP17/28 (Table S1) from 19 tracheophyte species, including 12 dicotyledons (*Brassica napus, Brassica oleracea, Brassica rapa, Beta vulgaris, Camelina sativa, Glycine max, Lotus japonicus, Medicago truncatula, Nicotiana tabacum, Pisum sativum, Solanum lycopersicum and Vitis vinifera*), 7 monocotyledons (*Ananas comosus, Asparagus officinalis, Hordeum vulgare subsp. vulgare, Musa acuminata, Oryza sativa, Triticum aestivum and Zea mays*) and 3 bryophytes (*Marchantia paleacea, Marchantia polymorpha and Physcomitrium patens*). Protein alignments confirmed that all identified sequences corresponded to bZIP proteins, showing the highly conserved bZIP domain composed by the basic region and the leucin loop region (Table S1, Figure S1A-B). The bZIP domain consists of two structural features located on a contiguous α- helix: a basic region of ∼16 amino acid residues containing a nuclear localization signal followed by an invariant N-x7-R/K motif that binds the DNA and a heptad repeat of leucines or other bulky hydrophobic amino acids (A/V/L/G/I/M/W/F/P) positioned exactly nine amino acids towards the C-terminus, creating an amphipathic helix (Jakoby et al., 2002; Zhang et al., 2016).

Contrary to numerous plants studied, which possess at least three bZIP involved in UPR signaling, only two grapevine sequences have been identified. The first protein sequence clustered within the dicot-specific clade of bZIP60 not far from the reference sequence of *A. thaliana* as shown in the phylogenetic tree (Figure 1A). Both *A. thaliana* and *V. vinifera* bZIP60 coding sequences share the same mRNA kissing stem-loop structures with the consensus sequences (CTGCTG) containing the putative splicing sites (Figure 1C). Based on the known splicing pattern described in *A. thaliana* (Iwata et al., 2025), we predicted a 23-nucleotides splicing which provokes a frameshift in the coding region, resulting in the synthesis of a new truncated protein. Structural prediction of the protein before and after splicing (Figure 1D) showed that splicing leads to the removal of the transmembrane domain (TMD) and the generation of an alternative open reading frame (ORF2), producing a new C-terminal region that includes a nuclear localization signal (NLS). These changes are consistent with the activation of bZIP60 via unconventional-splicing and its subsequent nuclear translocation, as described in *A. thaliana* (Nagashima et al., 2011). Additionally, the predicted 3D structure highlighted the conserved bZIP domain and transmembrane helix between *A. thaliana* and *V. vinifera* (Figure S2A), further supporting the identification of this sequence as VvbZIP60.

**Figure 1.**
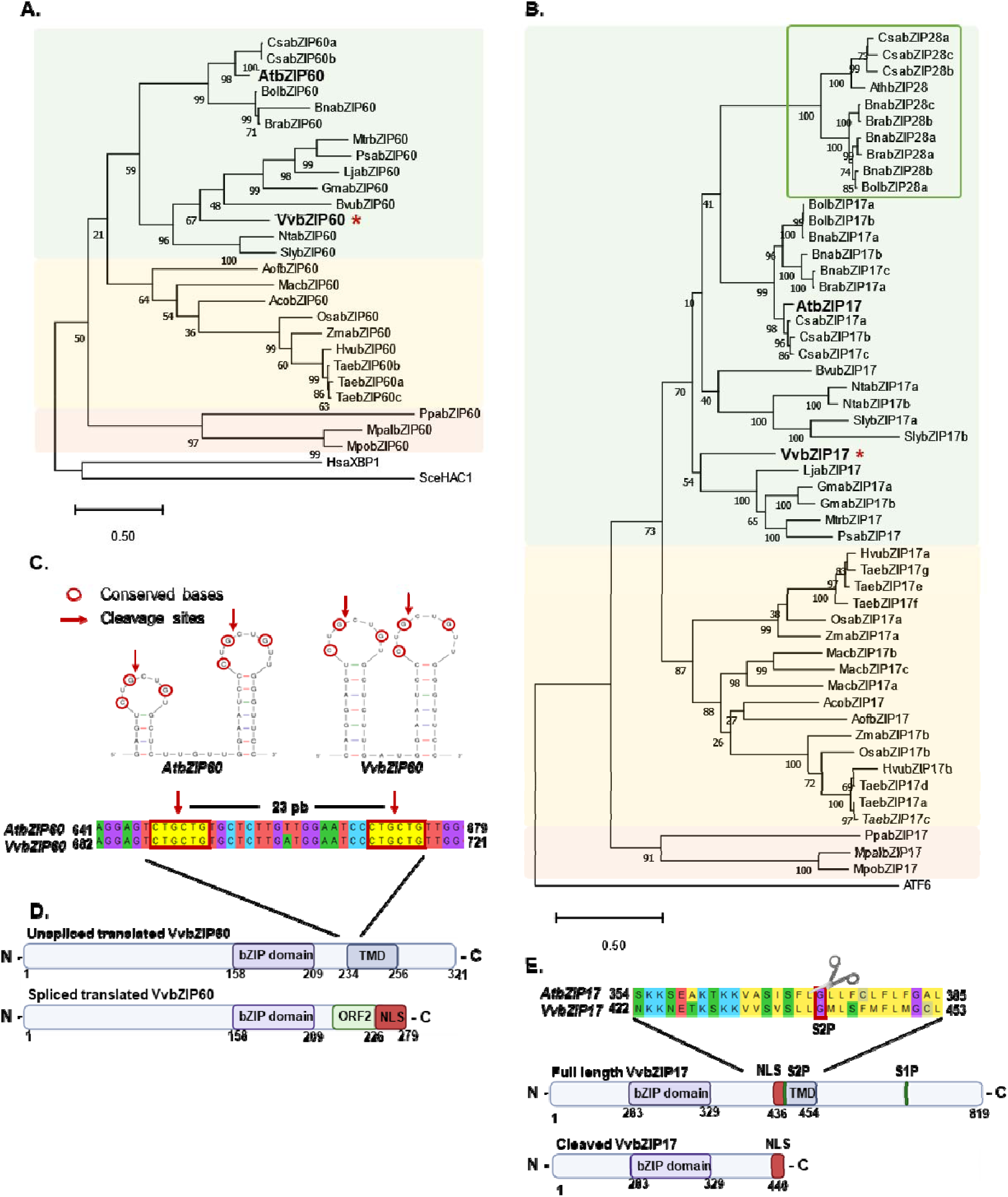
| Phylogenetic analysis of bZIP60 (A) and bZIP17 family (B). Phylogenetic tree of bryophyte (orange), angiosperms monocots (yellow) and eudicotyledons (green) represented in different colors. The phylogenetic tree is rooted with human (HsaXBP1) and yeast (SceHAC1) bZIP60 sequences (A) and human ATF6 sequence (B). Red asterisk indicated VvbZIP60 and VvbZIP17 sequences. The phylogenetic tree was constructed using MEGA11 software with the maximum-likelihood method and a bootstrap consensus of 1,000 bootstraps. **(C)** Twin hairpin loop structure at splicing site in *bZIP60* mRNA in *Arabidopsis thaliana* and *Vitis vinifera*. Each of the two loops contains three conserved bases (red rounds), also highlighted in yellow on the bZIP60 mRNA sequences. Red arrows indicate predicted cleavage sites. **(D)** Schematic representations of bZIP60 unspliced and spliced translated forms predicted *in silico*. In purple, the bZIP domain, in blue the transmembrane domain (TMD) predicted by “DAS - Transmembrane Prediction server” (Cserzö et all. 1997), in green the open reading frame generated by the frame shift (ORF2) predicted by “ORF Finder” (Stothard 2000) and in red the nuclear localization signal (NLS) predicted by “LOCALIZER” (Sperschneider J et al. 2017). **(E)** Schematic representations of bZIP17 before and after cleavage at S2P. In purple, the bZIP domain, in blue the transmembrane domain (TMD) predicted by “DAS - Transmembrane Prediction server” (Cserzö et all. 1997), in green the S2P and the S1P and in red the nuclear localization signal (NLS) predicted by “LOCALIZER” (Sperschneider J et al. 2017). The bZIP17 protein sequences from *Arabidopsis thaliana* and *Vitis vinifera* containing the conserved S2P cleavage site (scisors).

The second bZIP grapevine sequence grouped more closely to the bZIP17 sequences rather than within the clade including bZIP28 sequences, the latter resulted exclusively to the Brassicaceae family (Figure 1B). This finding suggests that in grapevine, as in most dicotyledons except for the Brassicaceae family, the only identified sequence is more closely related to bZIP17 than to bZIP28. Interestingly, within the monocots’ clade, two distinct subclades are observed, corresponding to the two putative rice bZIP17/28 sequences. Notably, OsbZIP39, here renamed OsbZIP17b, has been proposed to be the ortholog of Arabidopsis bZIP28 (Takahashi et al., 2012). In our analysis, OsbZIP17b does not cluster along with AtbZIP28, suggesting that monocots may possess two or more homologs of bZIP17 rather than both bZIP17 and bZIP28. For instance, in maize, ZmbZIP34, here named ZmbZIP17b, was first considered as the ortholog of AtbZIP28 (Zhu et al., 2019), while in later study, the same sequence have been referred as homolog of bZIP17, therefore naming it ZmbZIP17b (Ko and Brandizzi, 2021). In grapevine, the *in-silico* analysis of VvbZIP17 protein structure, showed a canonical basic leucine zipper (bZIP) domain near the N-terminus and a single transmembrane domain (TMD) positioned toward the C-terminus, consistent with its classification as a membrane-tethered transcription factor. The bZIP17 sequence alignment between *A. thaliana* and *V. vinifera* revealed also a conserved S1P motif cleavage (RRIL) in C-terminal and a S2P cleavage motif represented by a glycine within the hydrophobic TMD (Figure 1E, Table S1). Following proteolytic activation after ER stress, the N-terminal region is predicted to be cleaved, resulting in a truncated form of the protein retaining the bZIP domain but lacking the TMD (Figure 1E). Structural 3D alphafold prediction of the full-length VvbZIP17 superposed on AtbZIP17 protein structure highlighted the well-defined alpha-helical structures of the bZIP domain and the TMD (Figure S2B). These predictions support a conserved mechanism of activation through regulated intramembrane proteolysis under ER stress conditions in both plant species.

To complete the characterization of the UPR actors in grapevine, we chose to conduct a phylogenetic analysis also for the chaperone proteins BiP and the ER-transmembrane protein IRE1. As for bZIP17, bZIP28 and bZIP60, we used the reference sequences of *A. thaliana*. The BiP coding genes are known to be upregulated by the UPR pathway. In particular, *BiP3* is the most highly upregulated by abiotic stresses or by ER stress agents (Tajima et al., 2008; Herath et al., 2020) compared to *BiP1* and *BiP2*. As for *A. thaliana* we identified three grapevine sequences for BiP, two of them clustered within the BiP1/2 clade and the other one within the BiP3 sequences (Figure S3). We can also observe a third cluster separated from the BiP1/2 and BiP3 clades and specifically for the monocots. This group includes OsBiP4 and OsBiP5 which have been described to be involved in the ER stress response (Wakasa et al., 2012). To further characterize BiP1/2 and BiP3, we analyzed their promoter regions and searched for cis-regulatory elements (CREs), comparing them with the promoters of AtBiPs. Similar to Arabidopsis, several UPRE-III motifs were identified in the promoter regions of both VvBiP1/2, along with one ERSE-II and one UPRE-I motif (Figure S4A). In contrast, VvBiP3 contained only a single CRE, corresponding to the ERSE-I motif, which is also found exclusively in AtBiP3 (Figure S4B). These findings suggest that BiP1/2 and BiP3 possess distinct transcriptional regulatory binding sites and that VvbZIP TFs can bind to these ER stress-responsive cis-regulatory elements present in the promoters to upregulate UPR target genes such as VvBiPs.

Phylogenetic analysis of IRE1 sequences showed three distinct groups clustering each one with AtIRE1a, AtIRE1b and AtIRE1c respectively. The AtIRE1c clade did not root with the IRE1 sequences from human and yeast, in addition only Brassicaceae sequences were found, suggesting a specific and separate role in the Brassicaceae family for this protein. Two grapevine sequences were identified clustering each one with AtIRE1a and AtIRE1b respectively, suggesting two possible candidate orthologs for IRE1, here named VvIRE1a and VvIRE1b. (Figure S5). In addition, IRE1 proteins alignment showed the identification of the ATP binding motif (VAVKR) and the Ser/Thr protein kinase motif (DLKPQN) highly conserved among the IRE1a/b groups (Table S1). These results allowed us to identify the main putative UPR actors in grapevine, VvbZIP60 and VvbZIP17 among with VvBIPs and VvIRE1s compared to other plant species.

### 3.2. ER stress induces the splicing of *VvbZIP60* in grapevine cell suspension

The unconventional splicing of *bZIP60* mRNA has been shown previously in *A. thaliana* and other plant species (Moreno et al., 2012; Wang et al., 2017; Kaur and Kaitheri Kandoth, 2021; Li et al., 2022). To confirm this event in grapevine, we examined the splicing of *VvbZIP60* in grapevine cell suspension after 1 and 3 hours of DTT and TM treatments. We used the well-established PCR-based splicing assay with a set of primers flanking the splicing site (F-R primers, Figure 2A). *VvVPS54* transcript was used as a loading control. A single band of 248 pb corresponding to the unspliced form was observed in the control H_2_O- and DMSO-treated grapevine cell suspensions (Figure 2B). Interestingly, a second band of 225 pb, corresponding to the spliced forms of *VvbZIP60*, was detected in samples treated with DTT at both 1 and 3 h post treatment while TM treatments showed the second band only at 3 h (Figure 2B). A third band is also visible at around 250 pb, probably formed by the hybridization of bZIP60u and bZIP60s products during the PCR amplification. Such hybrid band has been previously observed and documented in RT-PCR analysis of *XBP-1* and *bZIP60* in Human and Arabidopsis, respectively (Shang and Lehrman, 2004; Moreno et al., 2012). Sequencing of the spliced and unspliced *VvbZIP60* PCR products confirmed the removal of a 23 bp fragment in the spliced form (Figure 2C). To further examine whether *Vitis vinifera bZIP60 mRNA* could undergo the unconventional splicing, the expression pattern of the spliced form was analyzed in grapevine cell suspension after 1 and 3 h of DTT and TM treatments. To specifically detects the spliced form of *VvbZIP60*, we designed a set of primers that specifically amplify the spliced and the unspliced versions. One of the reverse primers spans the two splice junction sites of the spliced *VvbZIP60* mRNA and can only anneal if the 23 bp have been removed during splicing, whereas the other anneals in the 23 bp of the unconventional intron (Figure 2A). qRT-PCR analysis confirmed the RT-PCR results: the spliced form of *VvbZIP60* was strongly induced by DTT at both 1 and 3 h, whereas TM strongly induced splicing only at 3 h (F-S primers, Figure 2D). These two independent methods confirm that *VvbZIP60* is regulated in a similar manner to other plant *bZIP60* and that the unconventional splicing happens at the predicted sites.

**Figure 2.**
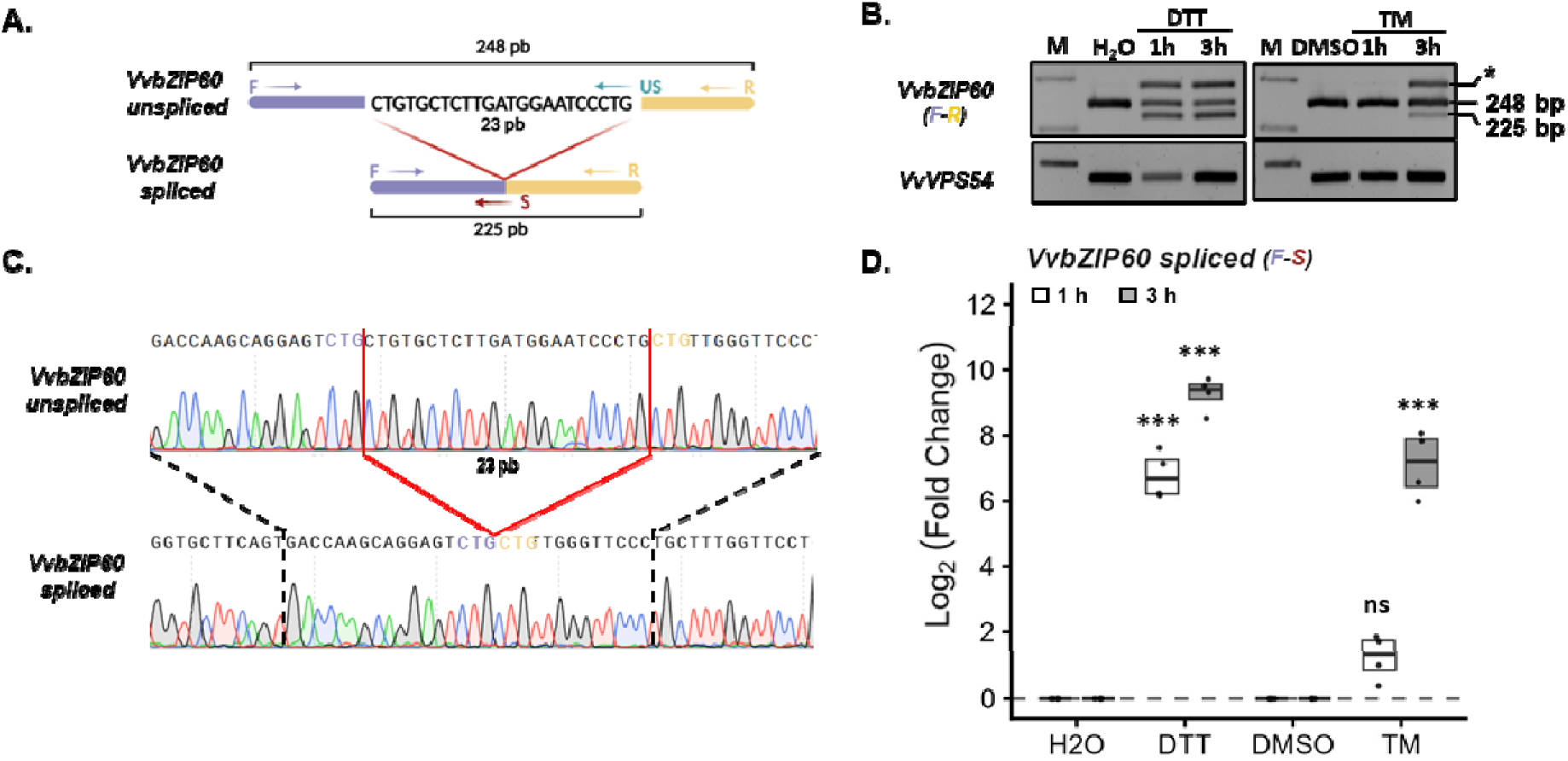
| Splicing assay of *bZIP60* in grapevine cell suspension. **(A)** Schematic representation of the primers used to amplify the spliced region of *VvbZIP60* for the qRT-PCR and the primers used for semi-quantitative RT-PCR. F-S primers for the putative splicing regions (red arrow) are indicated to amplify *bZIP60s* form. F-R primers are used to amplify the entire region covering the splicing site. **(B)** Semi-quantitative qRT-PCR analysis in agarose gel (3%) of *VvbZIP60* expression at 3 hours after 2 mM dithiothreitol (DTT), 0.2 μg/mL tunicamycin (TM) treatments and their respective negative controls with water or dimethyl sulfoxide (DMSO). *VvVPS54* was analyzed as a loading control. PCR product sizes are indicated at right. “M” indicated the DNA ladder. Asterisk indicates a hybrid band formed by the *bZIP60u* and *bZIP60s* PCR products. Such hybrid band has been also observed and documented in RT-PCR analysis of XBP-1 and bZIP60 in *A. thaliana* (Shang and Lehrman, 2004; Moreno et al., 2012). **(C)** Sanger sequencing chromatograms of *VvbZIP60 unspliced* (top) and *spliced* (bottom) forms. The red boxes highlight the 23-bp region present in the unspliced transcript, while the dashed lines mark the junction created by splicing and the consequent deletion of this fragment. **(D)** Log2-fold change values of *VvbZIP60 spliced* form measured by qRT-PCR at 1 and 3 h post treatment as in (B). Boxplots represent the distribution of four independent biological repeats (n=4). Means of technical duplicates (efficiency-weighted Cq(w) values) were normalized using mean Cq(w) data of two housekeeping genes (*VvVSP54 and VvRPL18B*) before being analyzed. Asterisks indicate statistically significant differences between negative controls (set at 0) and the relative treatments using ANOVA test followed by a Tukey’s HSD post hoc test, (P< 0.05).

### 3.3. IRE1-mediated *bZIP60* mRNA splicing is conserved in grapevine

In *A. thaliana*, IRE1 catalyzes the unconventional splicing of bZIP60 mRNA to produce the active transcription factor. In the double mutants *Atire1a/ire1b*, TM treatment failed to induce bZIP60 splicing conforming a central role of IRE1 in mediating the unconventional splicing (Nagashima et al., 2011; Zhang et al., 2016). After several failed attempts to obtain VvbZIPs CRISPR-Cas9 knock-out mutant, we used the IRE1-inhibitor STF-083010 to determine whether VvIRE1 mediates the unconventional splicing of bZIP60 mRNA in grapevine. In mammalian cells, STF-083010 binds the RNase domain of IRE1α and blocks cleavage of XBP1 mRNA without affecting the kinase activity (Chiu et al., 2023; Papandreou et al., 2011). We treated grapevine cell suspension with STF-083010 either together with the elicitor or 10 or 60 min before DTT or TM treatment, and we analyzed the expression of the main ER stress marker genes 1 h and 3 h after the proteotoxic treatment. In DTT-treated cells, co- or pre-treatment with STF-083010 reduced the level of *spliced VvbZIP60* at 1 h compared with DTT alone (Figure 3A). Surprisingly, this inhibitory effect was lost at 3 h, the pre-treated cells did not differ from cells treated with DTT alone. However, expression of *VvBiP3*, a well-known ER stress marker gene, was lower in the presence of the inhibitor at 3 h (Figure 3B). TM does not activate the UPR at 1 h, so the inhibitor had no visible effect at that time point. At 3 h, however, STF-083010 reduced both *VvbZIP60 splicing* and *VvBiP3* expression (Figure 3C-D). Interestingly, STF-083010 increased *VvIRE1b* expression at 3 h but not *VvIRE1a* (Figure S6). Together, these results support a role for *VvIRE1* in *VvbZIP60* mRNA-splicing in grapevine, suggesting that this mechanism is conserved in this species.

**Figure 3.**
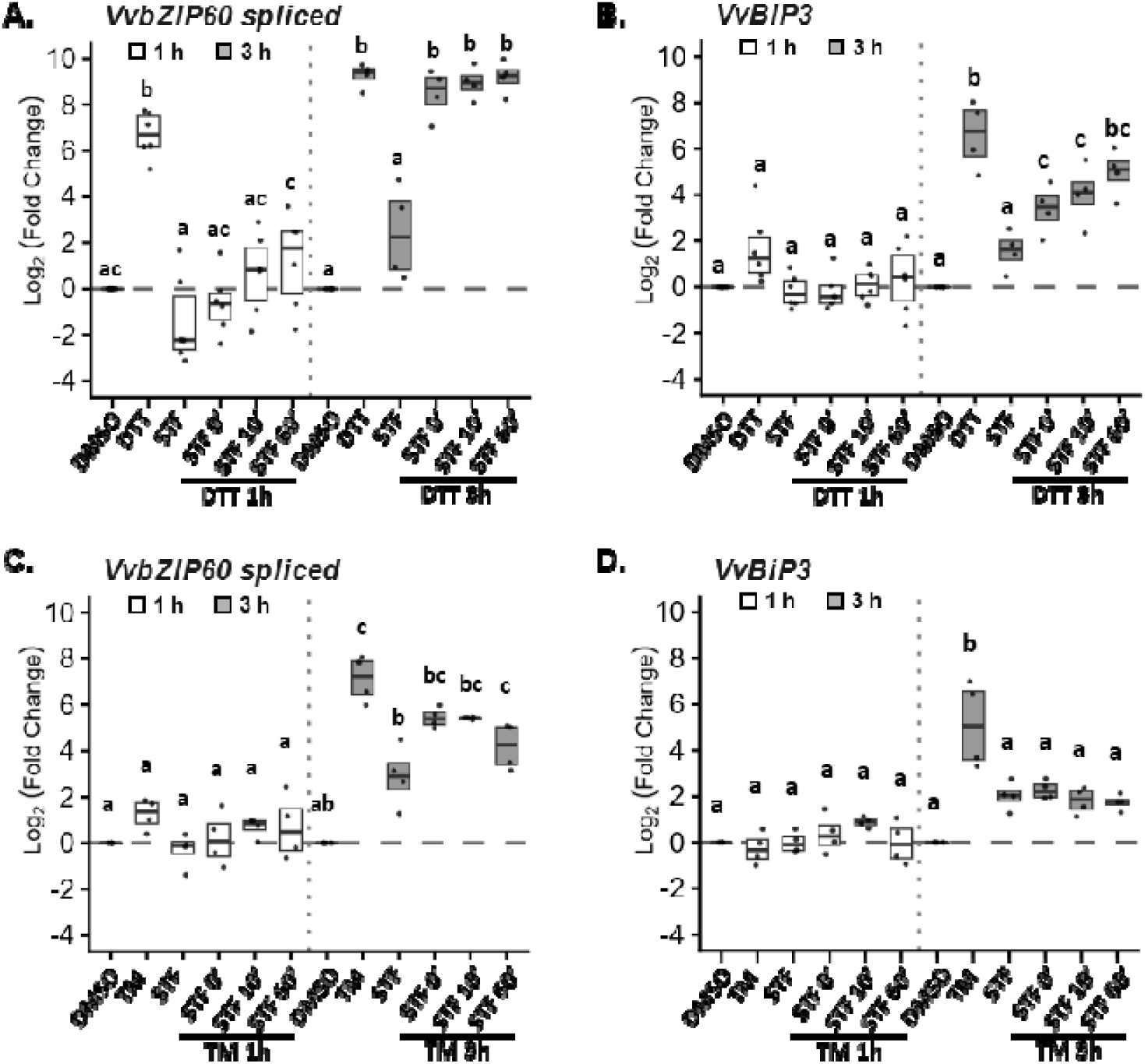
| The IRE1 inhibitor (STF-083010) decreases IRE1/bZIP60-related gene expression in grapevine. Log2-fold change values of VvbZIP60 spliced form and VvBiP3 measured by qRT-PCR after 1 h and 3 h of 2 mM dithiothreitol (DTT) (A-B) and 0.2 μg/mL tunicamycin (TM) (C-D) treatments following a pre-treatment of 0 - 10 and 60 min with 60μM STF-083010. Boxplots represent the distribution of at least four independent biological repeats (n≥4). Means of technical duplicates (efficiency-weighted Cq(w) values) were normalized using mean Cq(w) data of two housekeeping genes (VvVSP54 and VvRPL18B) before being analyzed. Letters indicate statistically significant differences among treatments for each time separately obtained using ANOVA test followed by a Tukey’s HSD post hoc test (P< 0.05). Dashed lines represent the negative control, DMSO (set at 0).

### 3.4. Heat stress activated the UPR and induces VvbZIPs re-localization to the nucleus

Previous results have confirmed the involvement of the UPR during different abiotic stresses such as heat, salt and drought (Liu et al., 2007a; Valente et al., 2009; Deng et al., 2011). It is well known that grapevine faces adverse environmental conditions all along its development, coping for example with extreme temperatures. In order to identify if heat stress could promote the activation of the UPR in grapevine, we incubated grapevine cell suspension at 37°C for 1 h and 3 h. In addition, DTT was also used as internal positive control. We chose to focus on the transcript response of three UPR genes: *VvbZIP60 spliced forms, VvbZIP17* and *VvBiP3*. As expected, DTT treatment strongly induced *VvbZIP60 splicing* and *VvBiP3* expression at both time points (Figure 4A). Similarly, heat stress (37°C) markedly induced *VvbZIP60 splicing*, with the strongest response at 1 h, as well as *VvbZIP17* and *VvBiP3* expression at both 1 h and 3 h (Figure 4A). To confirm whether VvbZIP17 and VvbZIP60 re-localize to the nucleus under heat-induced ER stress, mRFP-tagged versions of both transcription factors were transiently expressed in *Nicotiana benthamiana* leaves by agroinfiltration. Under control conditions, both RFP-tagged VvbZIP60 and VvbZIP17 localized to the ER, as shown by the merged image in Figure 4B. After 1 h at 37°C, however, both VvbZIPs re-localized from the ER to the nucleus, confirming the activation of VvbZIP60 unconventional splicing and of VvbZIP17 proteolytic cleavage in grapevine under ER stress. The presence of bZIP-binding cis-elements in the promoters of VvBIP genes (Figure S4), together with the stress-induced nuclear translocation of VvbZIP60 and VvbZIP17 (Figure 4B), supports a model in which these transcription factors directly activate *VvBIP* expression as part of the grapevine UPR signaling cascade. In addition to heat stress, a series of other abiotic stresses, including salt stress (NaCl), osmotic stress (PEG) and cold stress (4°C), were applied to grapevine cell suspension for 1 h and 3 h. Surprisingly, neither salt stress (NaCl) nor cold stress (4°C) upregulated UPR genes at either time point (Figure S7). In contrast, PEG-induced osmotic stress strongly upregulated the expression of *VvbZIP17* and *VvBiP3*, but only at 3 h, suggesting that osmotic stress differentially activates the two UPR arms, with the IRE1/bZIP60 and bZIP17 arms responding unevenly. Together, these results indicate that the UPR is activated by specific abiotic stresses, with differences in the timing of induction and in which arm of the pathway is engaged.

**Figure 4.**
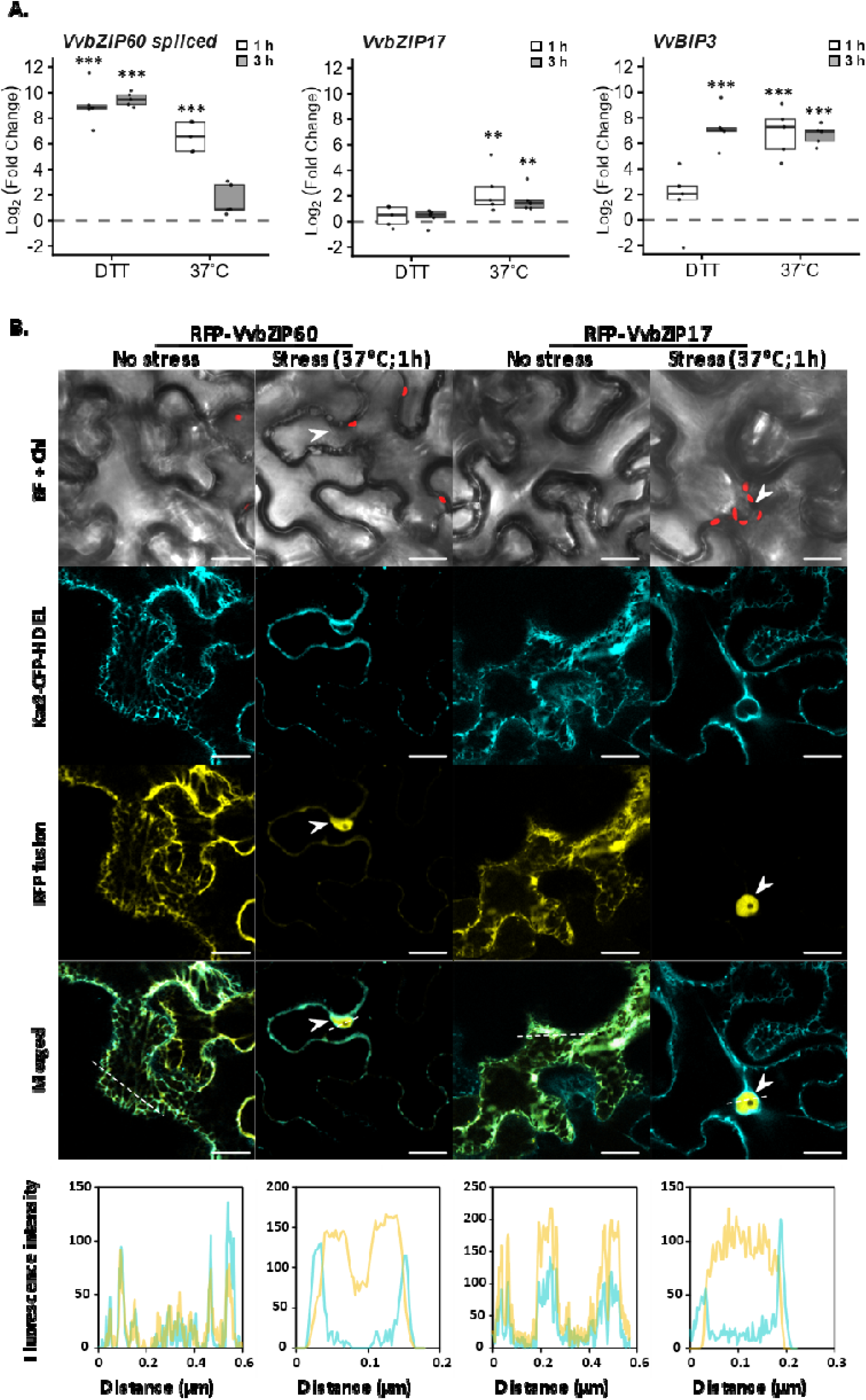
| Heat stress triggers UPR and induces the re-localization of both VvbZIP60 and VvbZIP17 to the nucleus. **(A)** Log2-fold change values of VvbZIP60 spliced form, VvbZIP17 and VvBiP3 measured by qRT-PCR 1 and 3 h after 2 mM dithiothreitol (DTT) or heat stress (37°C). Data represents the Log2-fold change of 4 independent biological repeats (n=4) normalized on the negative control (water) set at 0 (dot line). Means of technical duplicates (efficiency-weighted Cq(w) values) were normalized using mean Cq(w) data of two housekeeping genes (VvVSP54 and VvRPL18B) before being analyzed. Asterisks indicate statistically significant differences between negative control and the relative treatments using ANOVA test followed by a Tukey’s HSD post hoc test (P< 0.05). **(B)** Confocal microscopy images of N. benthamiana epidermal cells showing the subcellular localization of VvbZIP60 or VvbZIP17, 2 days post-infiltration, before and after 1 h at 37°C. Kar2-CFP-HDEL was used as ER marker. Merged images show colocalization of RFP- VvbZIP60 and RFP-VvbZIP17 with the ER marker. BF + Chl: transmitted light in bright field and chlorophyll autofluorescence. Arrows indicate the nucleus. Scale bars = 20 μm. The lower panels show fluorescence intensity profiles of CFP (cyan) and mRFP (yellow) along the white dashed line from the Merged panels above.

### 3.5. *VvbZIP17* and *VvbZIP60* gene expression are upregulated by chitosan

In previous works, bacterial microbe-associated molecular patterns (MAMPs) were shown to induce the expression of UPR genes (Arraño-Salinas et al., 2018; Chakraborty et al., 2020). Grapevine mainly faces several fungal pathogens and chitinaceous molecules derived from these interactions are known to trigger grapevine immunity (Brulé et al., 2019, 2024; Marzari et al., 2025b). To assess whether UPR is triggered by MAMPs, we treated grapevine cell suspensions with three different chitin oligomers (COs) of different degree of polymerization (DP): DP4 (CO4), DP6 (CO6) and chitosan DP6 (CHITO) and the most active epitope of the bacterial flagellin in grapevine (XcFlg22) (Trdá et al., 2014). We observed that none of these MAMP induced the UPR at 1 h and 3 h except for chitosan, which significantly increased the expression of all UPR genes at 3 h, except *VvBiP3* (Figure 5). Interestingly, chitosan was also the only MAMP to exhibit toxicity in cell suspensions: between 3 h and 24 h, the cell viability decreased by up to 50% compared to the control and other treatments (Figure S8). These results suggest a possible link between UPR and chitosan-induced cell death.

**Figure 5.**
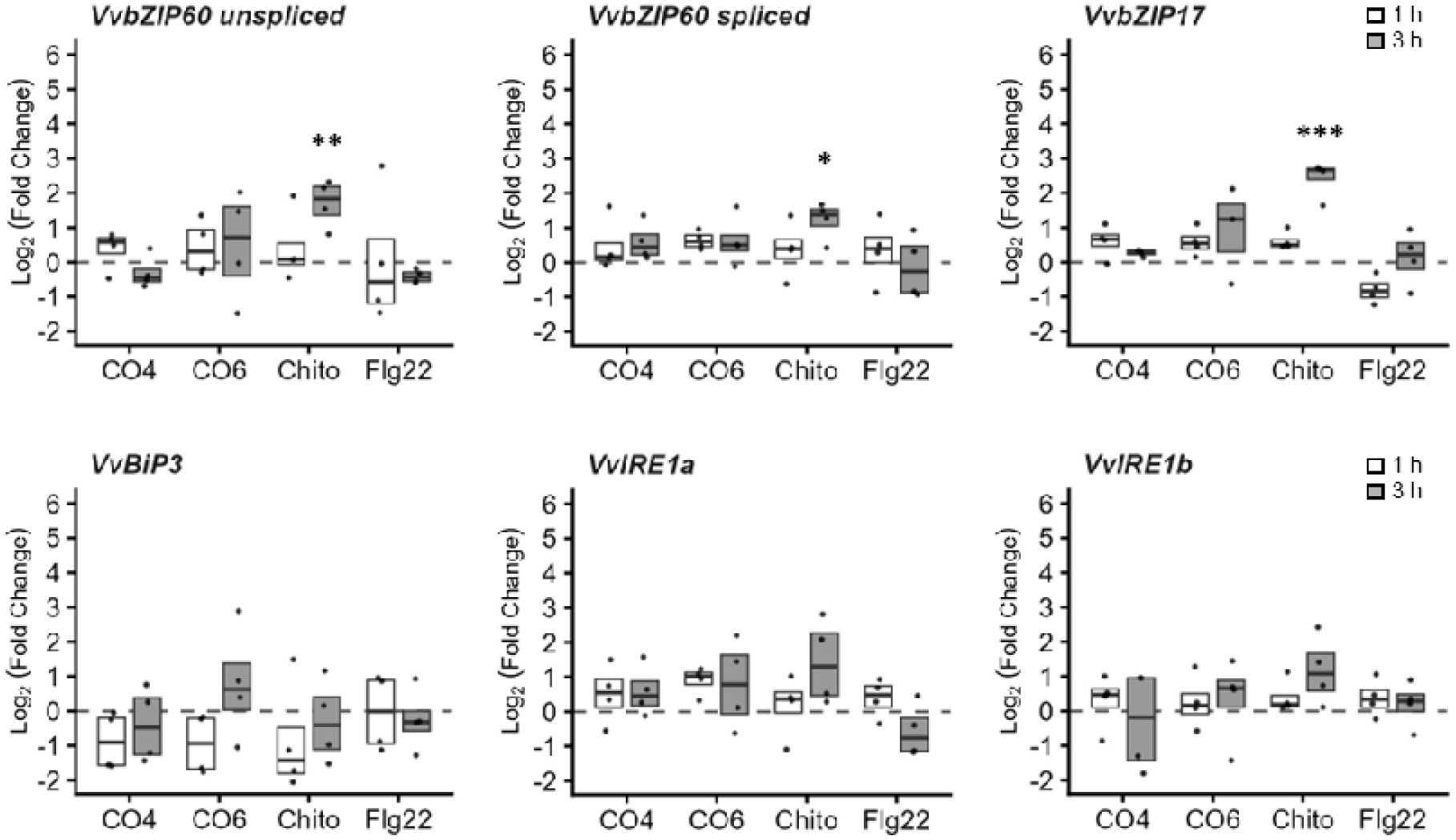
**| Chitosan triggers the expression of both UPR pathways**. Log2-fold change values of *VvbZIP60 unspliced* and *spliced* forms, *VvbZIP17, VvBiP3, VvIRE1a* and *VvIRE1b* measured by qRT-PCR 1 and 3 h after 0,2 mg/mL short-chitooligosaccharide DP4 (CO4), chitin DP6 (CO6), chitosan DP6 (CHITO) and 10^-8^ M Flg22. Data represents the Log2-fold change of four independent biological repeats (n=4) normalized on the negative control (water) set at 0 (dots line). Means of technical duplicates (efficiency-weighted Cq(w) values) were normalized using mean Cq(w) data of two housekeeping genes (*VvVSP54* and *VvRPL18B*) before being analyzed. Asterisks indicate significant differences (P < 0.05) between the negative control and the relative treatments. Normal data were analyzed via ANOVA with a Tukey’s HSD post hoc test; non-normal data were evaluated using a Kruskal-Wallis test with a Dunn’s post hoc test.

### 3.6. The grapevine pathogens *B. cinerea* and *P. viticola* trigger UPR pathways

Increased accumulation of UPR gene transcripts such as *bZIP60* and *bZIP17* have been reported during several plant-pathogen interactions (Xu et al., 2019; Blanchard et al., 2025). To investigate if grapevine pathogens influence the activation of the UPR, we inoculated grapevine cuttings grown in greenhouse with *P. viticola*, the causal agent of downy mildew and *B. cinerea* which is responsible for the grey mould disease. *P. viticola* was applied directly to the leaves of whole grapevine cuttings (Figure S9A), whereas *B. cinerea* was inoculated on leaf disks (Figure S9C). Infection was confirmed by quantifying transcript levels of grapevine immune genes, such as acidic chitinase (*CHIT4C*) and stilbene synthase (*STS*), using qRT-PCR at various time points. Compared to the water-treated control plants, both pathogens strongly induced the defense-related genes at all time points (Figure S9B, D). At 5 days post inoculation (dpi), *P. viticola* triggered the splicing of *VvbZIP60* and increased the expression level of *VvbZIP17*, *VvBiP3* and *VvIRE1b* (Figure 6A). At 7 dpi, only the *VvbZIP60* spliced form and *VvIRE1b* were significantly upregulated (Figure 6A). At 5 dpi, *P. viticola* did not reach full sporulation since mycelium was not visible, whereas at 7 dpi the abaxial side of the leaves was fully colonized by the oomycete (Figure S9A).

**Figure 6.**
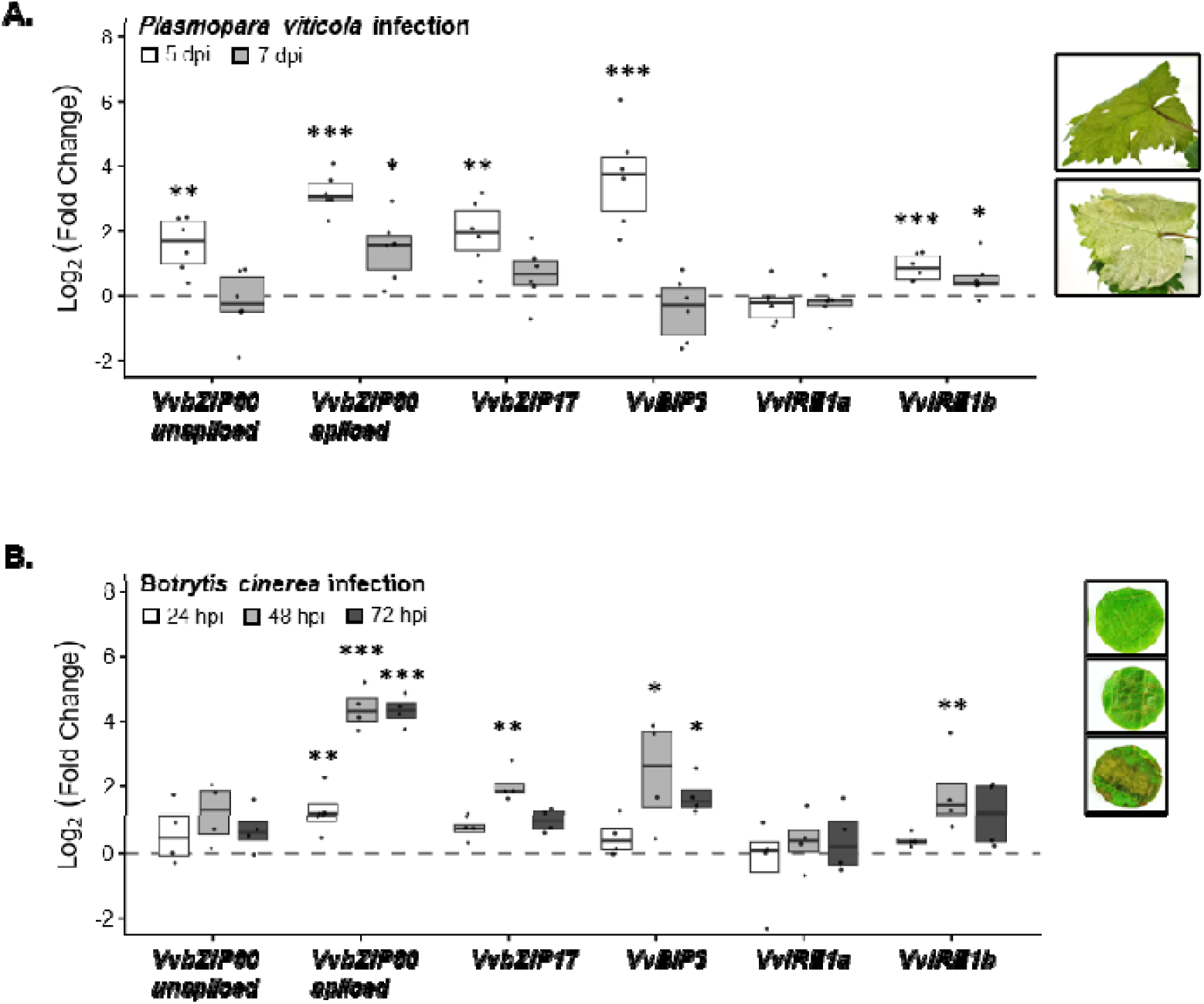
| P. viticola and B. cinerea trigger the expression of both UPR pathways. **(A)** Log2- fold change values of VvbZIP60 unspliced and spliced forms, VvbZIP17, VvBiP3, VvIRE1a and VvIRE1b measured by qRT-PCR 5-7 days post infection (dpi) with P. viticola on grapevine leaves **(B)** Log2-fold change values of VvbZIP60 unspliced and spliced form, VvbZIP17, VvBIP3, VvIRE1a and VvIRE1b measured by qRT-PCR at 24, 48 or 72 hours post infection (hpi) with B. cinerea on grapevine leaf disks. Boxplots represent the distribution of six and four independent biological repeats (n=6/n=4) respectively. Means of technical duplicates (efficiency-weighted Cq(w) values) were normalized using mean Cq(w) data of two housekeeping genes (VvVATP16 and VvEF1a) before being analyzed. Asterisks indicate significant differences (P < 0.05) between the negative control and the relative treatments. Normal data were analyzed via ANOVA with a Tukey’s HSD post hoc test; non-normal data were evaluated using a Kruskal-Wallis test with a Dunn’s post hoc test.

In this context, it is interesting to note that despite the *VvbZIP60* splicing we do not observe *VvBiP3* upregulation at 7 dpi. These results suggest that ER stress might be an early response during *P. viticola* infection. Similarly, *B. cinerea* also induced the activation of all UPR genes in grapevine leaves at 24, 48 or 72 h post inoculation (hpi) with a strong induction of the *VvbZIP60* splicing at all time points (Figure 6B). *VvbZIP17* and *VvIRE1b* showed an increased expression only at 48 hpi, while *VvBiP3* was significantly induced at 48 and 72 hpi (Fig.6B). These results show that UPR is involved in responses to *B. cinerea* infection, from early time points to necrotic symptoms in grapevine.

### 3.7. Inhibition of IRE1-mediated VvbZIP60 unconventional splicing increases *B. cinerea* infection

Since several recent studies have demonstrated an interplay between ER proteostasis and plant immunity (Marzari et al., 2025a; Meng et al., 2026), we wondered whether inhibition of IRE1 RNase activity could play an active role during direct interaction with a pathogen. To test this hypothesis, we conducted pathogenicity assays in which leaf discs pre-treated with STF-083010 for 1 h were inoculated with either *B. cinerea* or *P. viticola*. Disease development was monitored at 3 or 7 dpi, respectively. Leaf discs pre-treated with STF-083010 and challenged with *Plasmopara viticola* showed 20% lower sporulation (Figure S10A). However, because proteostasis may contribute to germination and pathogenicity in phytopathogens such as oomycetes (Ai et al., 2026), we performed a zoospore encystment assay on STF-083010 pre-treated leaf discs. The number of encysted zoospores differed markedly between pre-treated and control discs (Figure S10B-C), suggesting that STF-083010 affects the ability of *P. viticola* zoospores to settle on the leaf surface. The reduced sporulation may therefore reflect a direct effect of STF-083010 on *P. viticola* rather than on the plant UPR, and we cannot conclude that the UPR is involved in downy mildew infection.

Regarding *Botrytis cinerea* infection, pre-treated leaf discs showed a larger necrotic area than controls. On average, necrotic area reached 46.04 mm² in the pre-treated condition versus 37.93 mm² in the control, corresponding to a 21% increase (Figure 7). To determine whether this effect was due to IRE1 inhibition rather than a direct effect on the pathogen, we performed a *B. cinerea* growth assay, which showed that STF-083010 had no effect on fungal development (Figure S11). Collectively, these results indicate that IRE1 ribonuclease activity contributes to grapevine resistance to grey mold disease.

**Figure 7.**
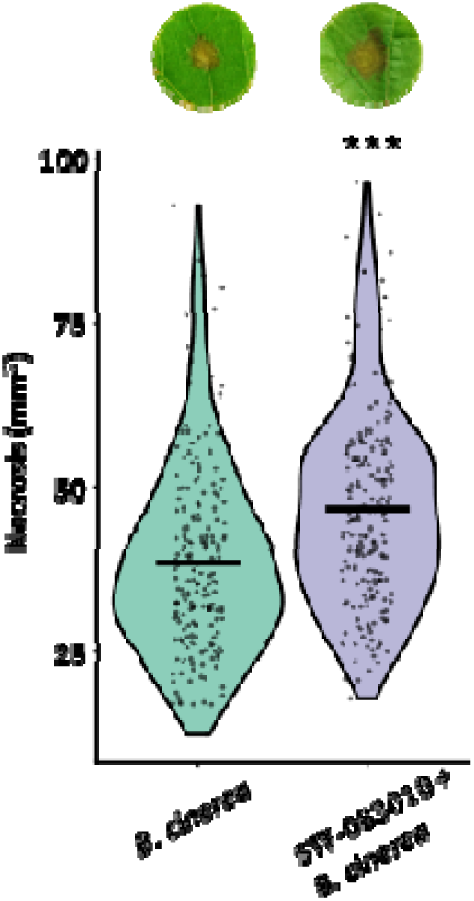
**| The IRE1 inhibitor (STF-083010) increases *B. cinerea* infection but does not affect its viability**. Necrotic lesion areas (mm²) were measured on individual leaf discs and compared between STF-083010-treated (60μM) and H₂O-treated conditions using ImageJ. For each biological replicate and condition, necrotic area was quantified on approximately 60 leaf discs; the mean value per biological replicate was then used as the statistical unit (n = 3 independent biological replicates). A generalized linear mixed model (GLMM) with a Gamma distribution and a log link function, with treatment included as a fixed effect and biological replicate as a random effect (random intercept) was used to analyze the necrosis. Black line represents the mean p<0,001.

## 4. Discussion

An increasing number of studies have demonstrated a strong link between the plant’s surrounding environment, including both abiotic factors and microorganisms’ interaction, and the activation of the UPR (Ko and Brandizzi, 2024; Adhikari et al., 2025; Liu et al., 2025; Marzari et al., 2025a; Meng et al., 2026). In this study, we identified the grapevine ortholog of bZIP60, designated as VvbZIP60 and demonstrated that its transcript undergoes unconventional splicing in response to ER stress induced by DTT and TM (Figure 2) and that this mechanism is mediated by VvIRE1 as shown in the IRE1-inhibition assay (Figure 3). Notably, VvbZIP60 harbors two conserved kissing stem-loop structures in its mRNA, essential for IRE1-mediated splicing. We also identified two putative grapevine orthologs of the ER-transmembrane sensor IRE1, VvIRE1a and VvIRE1b. Interestingly, *VvIRE1b* consistently exhibited higher expression levels than *VvIRE1a* under most of the stress conditions analyzed, notably after IRE1 inhibitor STF-083010 treatment (Figure S6), suggesting a potentially more prominent role in initiating the UPR in grapevine. Although AtIRE1a and AtIRE1b are both able to splice AtbZIP60, and the mutants *Atire1a* and *Atire1b* exhibit the same phenotype under ER stress induced by TM, AtIRE1b plays a dominant role in *bZIP60* splicing (Deng et al., 2011; Moreno et al., 2012), suggesting a partial functional redundancy (Nagashima et al., 2011). Interestingly, *VvIRE1b* is also more highly induced upon pathogen infection and notably *B. cinerea* (Figure 6A-B) in contrast to *A. thaliana*, in which neither *AtIRE1a* nor *AtIRE1b* is transcriptionally induced upon infection (Blanchard et al., 2025). In addition, AtIRE1b’s RNase activity is required for ER stress-induced autophagy via RIDD, independently of bZIP60 (Bao et al., 2018). Given that autophagy contributes to plant defense against necrotrophic fungi like *B. cinerea* (Lai et al., 2011), VvIRE1b may also induce RIDD-autophagy pathways in grapevine stress responses and thus participate in grapevine defense against *B. cinerea* in a different way than AtIRE1. These results indicate that although VvIRE1 and AtIRE1 isoforms are close orthologs, further studies are needed in grapevine to clarify the functional roles of both IRE1 isoforms, in the context of *bZIP60* mRNA splicing, RIDD-induced autophagy, and defense against necrotrophic fungi. The second arm of the UPR in plants is defined by the bZIP17/28 signaling module. Interestingly, our phylogenetic analysis revealed the presence of a single grapevine bZIP sequence clustering with bZIP17, while no grapevine sequence grouped with the bZIP28 clade. In the dicots, bZIP28 orthologs were found to be exclusive members of the Brassicaceae. This arm of the pathway may be therefore lineage-specific and absent in non-Brassicaceae species such as *Vitis vinifera*. This finding highlights possible evolutionary divergence in UPR signaling components among dicotyledonous species and raises questions about how UPR is regulated in species showing only one bZIP17/28 ortholog, whereas stress responses are usually shared between two transcription factors in plants already analyzed.

The UPR is known to be induced by different abiotic and biotic stresses. Here, we observed a significant induction of both UPR arms, under a variety of stress conditions. Among the abiotic factors tested in this study, heat and osmotic stress were the strongest ER-stressors. Heat stress is a well-known ER-stress inducer which triggers the splicing of *bZIP60* in different plant species such as *A. thaliana*, tomato and rice (Deng et al., 2011; Takahashi et al., 2012; Kaur and Kaitheri Kandoth, 2021). On the other hand, bZIP17 was only described to play important roles in maintaining fertility under heat stress conditions in *Arabidopsis* (Gao et al., 2022). Here, we showed a high induction of *VvbZIP60 splicing* and *VvbZIP17* transcript levels after 1 h at 37°C (Figure 4A) but also that heat stress induces the re-localization of both VvbZIP60 and VvbZIP17 from the ER to the nucleus (Figure 4B). Nuclear re-localization of bZIP17 has previously been shown under salt stress in *Arabidopsis* (Liu et al., 2007a), and bZIP60 re-localization has been shown under DTT treatment (Parra-Rojas et al., 2015); however, to our knowledge, this is the first report of heat-induced re-localization of both transcription factors in a single system, and the first in grapevine. Alongside the induction of *VvbZIP60* splicing and VvbZIP17, *VvBiP3* was also strongly upregulated. Interestingly, it has been shown that the activation of *AtBIP3* by *AtbZIP60* depends on the ERSE-I cis-elements in its promoter (Iwata and Koizumi, 2005b). Notably, we identified this motif exclusively in the promoter region of *VvBiP3*, suggesting that VvbZIP60 may specifically trigger its expression. Interestingly, in our work we demonstrated that PEG-induced osmotic stress also triggers the UPR but but activates only the bZIP17 arm (Figure S7). A previous transcriptional analysis revealed that *VvbZIP17* was upregulated in responses to drought stress (Liu et al., 2014) but no previous study has focused on PEG-induced osmotic stress specifically. Interestingly, in *A. thaliana*, bZIP17 was found to be more involved in the response to salt (Liu et al., 2007a). In grapevine cell suspension, NaCl treatment did not induce the UPR. Grapevine is relatively tolerant to salt stress, likely due to its adaptation to drought-prone environments. In a recent study, a transcriptome analysis of all *bZIP* TFs was conducted in grapevine to analysis their response to salt stress. Several *bZIPs* were upregulated under salt stress however, neither *VvbZIP17* nor *VvbZIP60* showed significant changes in expression (Guan et al., 2018; Çakır Aydemir et al., 2020). These results highlight the complexity of the UPR and suggest that its activation may vary between plant species.

The primary recognition of microorganisms involves the perception of highly conserved molecular signatures, also known as MAMPs. MAMPs are found in many types of microorganisms, including bacteria, fungi, viruses and protozoa and tend to be highly conserved, meaning that they are structurally similar across different species of microbes (Newman et al., 2013). A study carried out in *A. thaliana* seedlings showed that the UPR genes, notably *IRE1a* and *IRE1b*, were induced after treatment with the flagellin-derived peptide flg22 and elongation factor Tu-derived peptide elf18, suggesting an impact of these MAMPs on the ER-homeostasis (Chakraborty et al., 2020). Here, we tested several MAMPs, including flg22 and chitinous molecules such as chitin of different DPs and chitosan. Surprisingly, only chitosan could induce the UPR, particularly the transcription of *VvbZIP17* (Figure 5). Chitosan is a deacetylated derivative of chitin and is known to be a potent elicitor of plant immunity. In grapevine it exhibits antimicrobial properties against *B. cinerea*, *P. viticola* and Esca disease agents (Mian et al., 2023; Martín et al., 2023; Brulé et al., 2024). Chitosan-triggered immunity is associated with a set of cellular and molecular changes including cell wall lignification and callose deposition, ROS production, ion fluxes associated with the plasma membrane depolarization, MAPK activation, biosynthesis of phytoalexins and defense-responsive proteins (El Hadrami et al., 2010; Narula et al., 2020; Brulé et al., 2024). Interestingly, it has been demonstrated that ROS production affects the ER stress response in plants, acting as secondary messengers (Ozgur et al., 2018) and that UPR is also involved in lignin deposition (Nakamura et al., 2022). In addition, chitosan has been shown to trigger cell death in grapevine cell suspension (this study) and in soybean cells, where it occurs via activation of a caspase-3-like protease (Zuppini et al., 2004). This protease activity is well-known to be involved in ER stress-induced plant cell death (Cai et al., 2014). These findings suggest that chitosan-induced cell death may be linked to UPR signaling through protease activity, although further studies are needed to confirm this connection.

In this study, we showed that *P. viticola* and *B. cinerea* upregulate all UPR genes with the exception of *VvIRE1a* in grapevine leaves (Figure 7), suggesting an interplay between plant immunity and proteostasis. In addition, we demonstrated that inhibition of IRE1-mediated bZIP60 mRNA unconventional splicing directly affect grapevine susceptibility to *B. cinerea*, increasing pathogen infection up to 20% (Figure 8). These results are similar to previous studies which showed the importance of a functional IRE1-bZIP60 pathway for pathogen resistance (Blanchard et al., 2025; Moreno et al., 2012). In particular, Blanchard et al. demonstrated that double mutant *ire1a-ire1b* in *A. thaliana* was more susceptible to the infection, showing a 17% increase in lesion diameter relative to the WT genotype (Blanchard et al., 2025). Interestingly, STF-083010 pre-treatment had no effect on *B. cinerea* growth *in vitro* (Figure S11), whereas it strongly reduced the number of viable *P. viticola* zoospores on treated leaf discs (Figure S10B-C). It is possible that, since STF-083010 is a general IRE1 inhibitor, it may also affect the microorganism’s own UPR pathway. Several studies have already shown that perturbation of a pathogen’s UPR can influence the outcome of infection (Marzari et al., 2025a). For instance, in *Candida albicans*, an *ire1*-mutant strain shows pleiotropic defects in ER stress response, antifungal tolerance, cell wall regulation and virulence-related traits (Sircaik et al., 2021). Similarly, in the fungal pathogen *Aspergillus fumigatus*, STF-083010 inhibition of the IRE1 ortholog blocks the unconventional splicing of the bZIP60 ortholog *hacA* and sensitized the fungus to carvacrol-induced ER stress (Guirao-Abad et al., 2020). However, it is worth noting that *P. viticola* zoospores, unlike *B. cinerea* conidia, lack a protective cell wall and rely solely on their plasma membrane for structural integrity (Hardham, 2007). It is therefore also possible that STF-083010 exerts a direct effect on this particularly fragile cell type. Finally, to our knowledge, no study has yet investigated the canonical UPR mechanisms in oomycetes, and this would be a valuable direction for future research to clarify whether the effect of STF-083010 on *P. viticola* reflects inhibition of a pathogen-intrinsic UPR, a direct effect, or both.

To better understand the specific role of each UPR component, more in-depth functional genomic studies will be required. Recent findings show that the *bzip17* single mutant is more resistant to *B. cinerea* infection, suggesting that bZIP17 may also act as a negative regulator of defense (Blanchard et al., 2025). Silencing *bZIP17* in grapevine by RNAi or Virus-Induced Gene Silencing (VIGS) would help determine whether this function is conserved in this species. In addition, multi-omic approaches combining transcriptomics, proteomics and ChIP-seq would allow the fine-tuned regulation of the UPR to be dissected, by identifying the target genes of each transcription factor and their dynamics during infection, as already done in *Arabidopsis* (Ko and Brandizzi, 2022a). Finally, genome-wide association studies (GWAS) in grapevine could reveal natural genetic variation in UPR genes associated with resistance to different pathogens.

## Acknowledgements

We acknowledge Dr. Benoit Darblade (Elicityl, Crolles, France) for the CO6, CO4 and chitosan oligomers. Subcellular localizations were performed at the DImaCell Facility. The authors thank Chrystel DEULVOT for help with microscopy data acquisition.

## CRediT authorship contribution statement

**Tania Marzari:** Writing-original draft, Investigation, Formal analysis, Data Curation. Visualisation. Conceptualization. **Emma Poilvert:** Methodology, Investigation. **Cécile Blanchard:** Investigation. **Karine Palavioux:** Resources. **Agnès Klinguer:** Resources. **Hugo Hassangatebois:** Resources. **Alix Martinet:** Investigation. **Aurélien Henry:** Investigation. **Benoit Poinssot:** Writing-review & editing, Supervision, Project administration. Funding acquisition, Conceptualization. **Mathieu Gayral:** Writing-review & editing, Supervision, Project administration. Funding acquisition, Conceptualization.

## Data availability

All data supporting the findings of this study are available within the paper and its supporting information are published online, the materials of this study are available from the corresponding author upon reasonable request.

## Declaration of generative AI and AI-assisted technologies in the writing process

During the preparation of this work the authors used “openAI” in the writing process to improve the readability and language of the manuscript. After using this tool, the authors reviewed and edited the content as needed and take full responsibility for the content of the published article.

