## Supplementary Figures for "Abiotic and biotic stress activate the unfolded protein response in grapevine, with IRE1/bZIP60 signaling contributing to *Botrytis cinerea* resistance"

### Slide 1
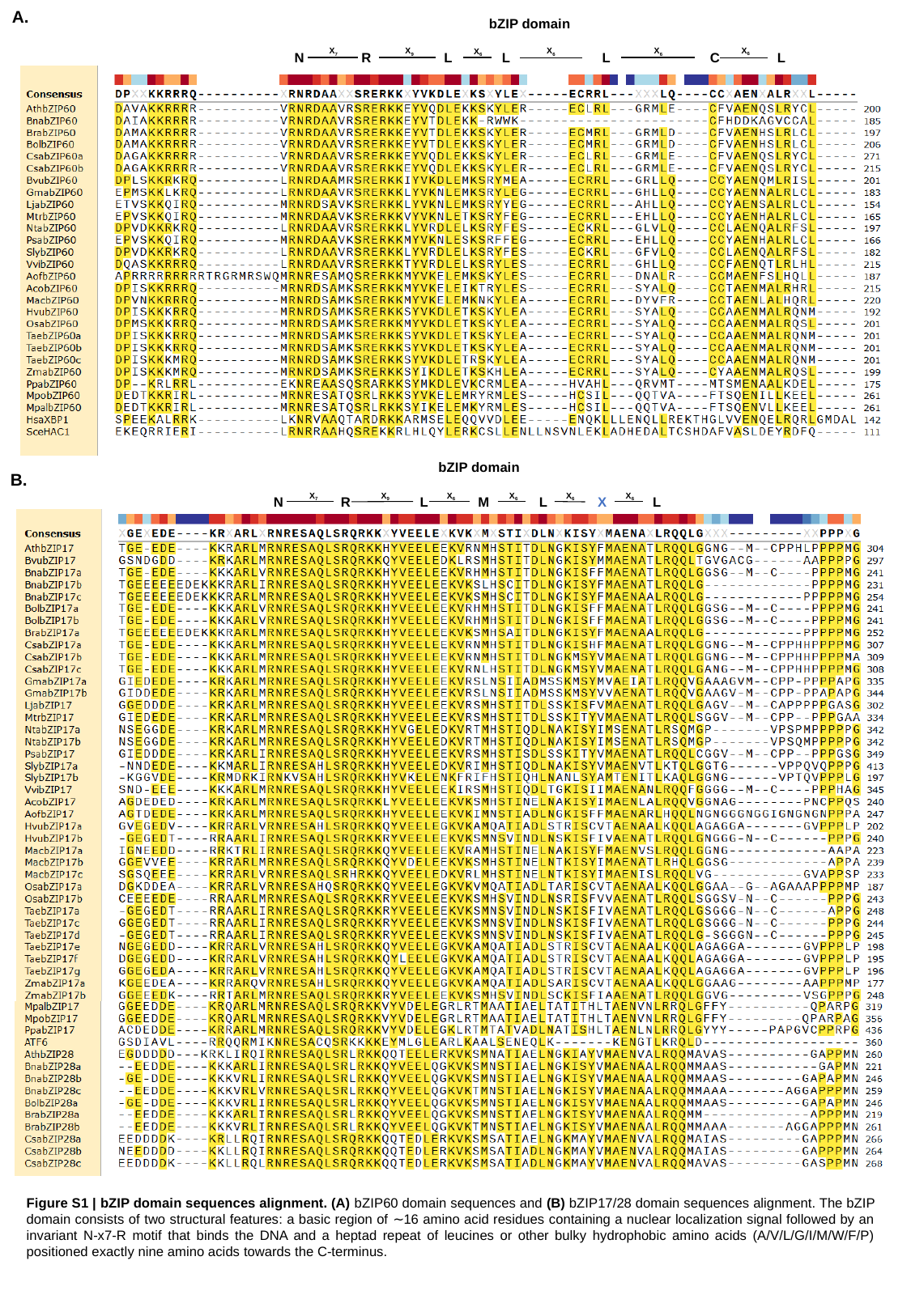

A.
bZIP domain
X_6_
X_6_
X_6_
X_7_
X_9_
X_6_
L L L C L
N R
bZIP domain
X_6_
X_6_
X_6_
X_6_
X_7_
X_9_
N R
 L M L X L
B.
Figure S1 | bZIP domain sequences alignment. (A) bZIP60 domain sequences and (B) bZIP17/28 domain sequences alignment. The bZIP domain consists of two structural features: a basic region of ∼16 amino acid residues containing a nuclear localization signal followed by an invariant N-x7-R motif that binds the DNA and a heptad repeat of leucines or other bulky hydrophobic amino acids (A/V/L/G/I/M/W/F/P) positioned exactly nine amino acids towards the C-terminus.

### Slide 2
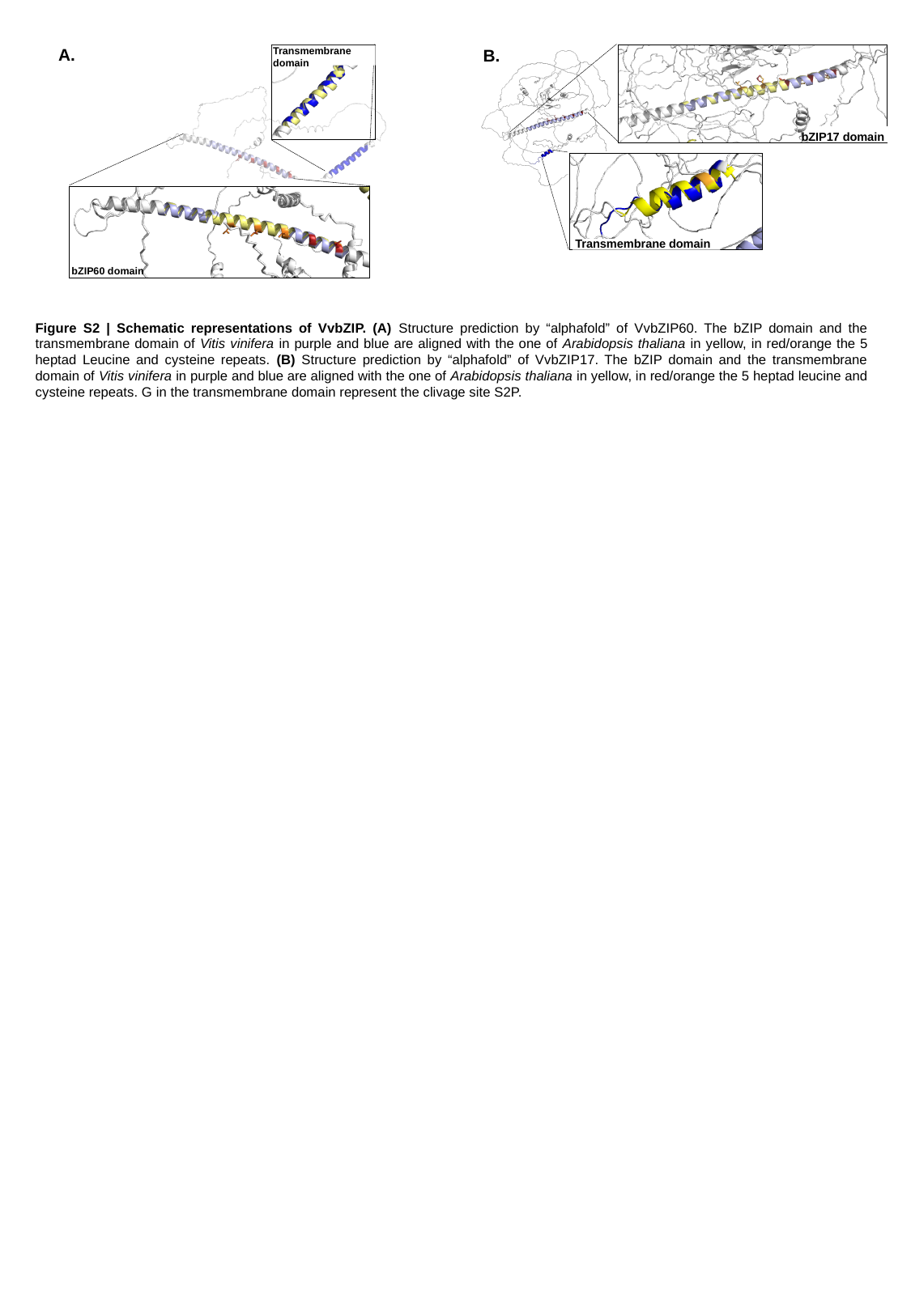

A.
Transmembrane
domain
bZIP60 domain
B.
bZIP17 domain
Transmembrane domain
Figure S2 | Schematic representations of VvbZIP. (A) Structure prediction by “alphafold” of VvbZIP60. The bZIP domain and the transmembrane domain of Vitis vinifera in purple and blue are aligned with the one of Arabidopsis thaliana in yellow, in red/orange the 5 heptad Leucine and cysteine repeats. (B) Structure prediction by “alphafold” of VvbZIP17. The bZIP domain and the transmembrane domain of Vitis vinifera in purple and blue are aligned with the one of Arabidopsis thaliana in yellow, in red/orange the 5 heptad leucine and cysteine repeats. G in the transmembrane domain represent the clivage site S2P.

### Slide 3
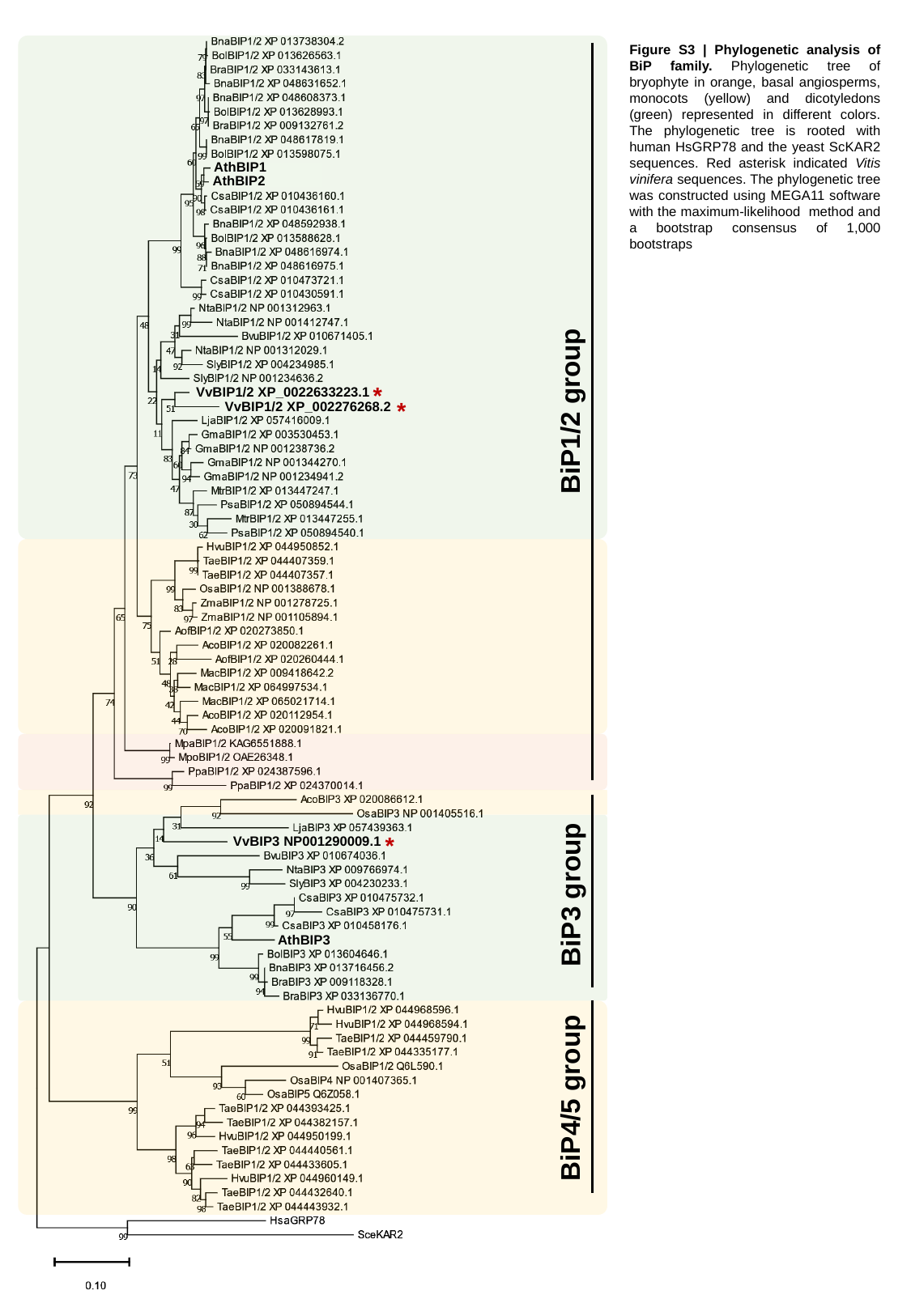

AthBIP1
AthBIP2
*
BiP1/2 group
*
*
BiP3 group
AthBIP3
BiP4/5 group
Figure S3 | Phylogenetic analysis of BiP family. Phylogenetic tree of bryophyte in orange, basal angiosperms, monocots (yellow) and dicotyledons (green) represented in different colors. The phylogenetic tree is rooted with human HsGRP78 and the yeast ScKAR2 sequences. Red asterisk indicated Vitis vinifera sequences. The phylogenetic tree was constructed using MEGA11 software with the maximum-likelihood method and a bootstrap consensus of 1,000 bootstraps
VvBIP1/2 XP_0022633223.1
VvBIP1/2 XP_002276268.2
VvBIP3 NP001290009.1

### Slide 4
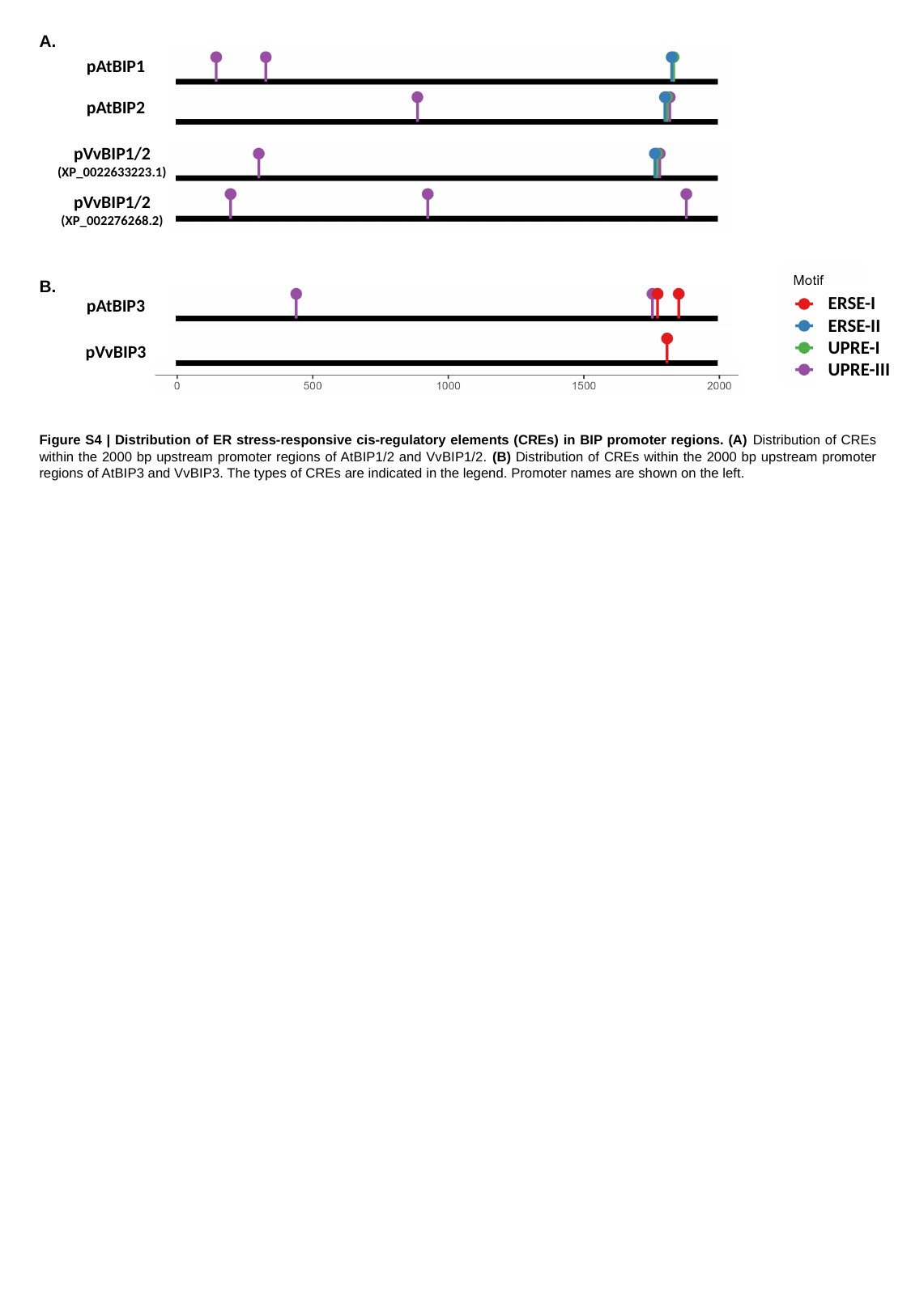

A.
pAtBIP1
pAtBIP2
pVvBIP1/2
(XP_0022633223.1)
pVvBIP1/2
(XP_002276268.2)
pAtBIP3
pVvBIP3
B.
ERSE-I
ERSE-II
UPRE-I
UPRE-III
Figure S4 | Distribution of ER stress-responsive cis-regulatory elements (CREs) in BIP promoter regions. (A) Distribution of CREs within the 2000 bp upstream promoter regions of AtBIP1/2 and VvBIP1/2. (B) Distribution of CREs within the 2000 bp upstream promoter regions of AtBIP3 and VvBIP3. The types of CREs are indicated in the legend. Promoter names are shown on the left.

### Slide 5
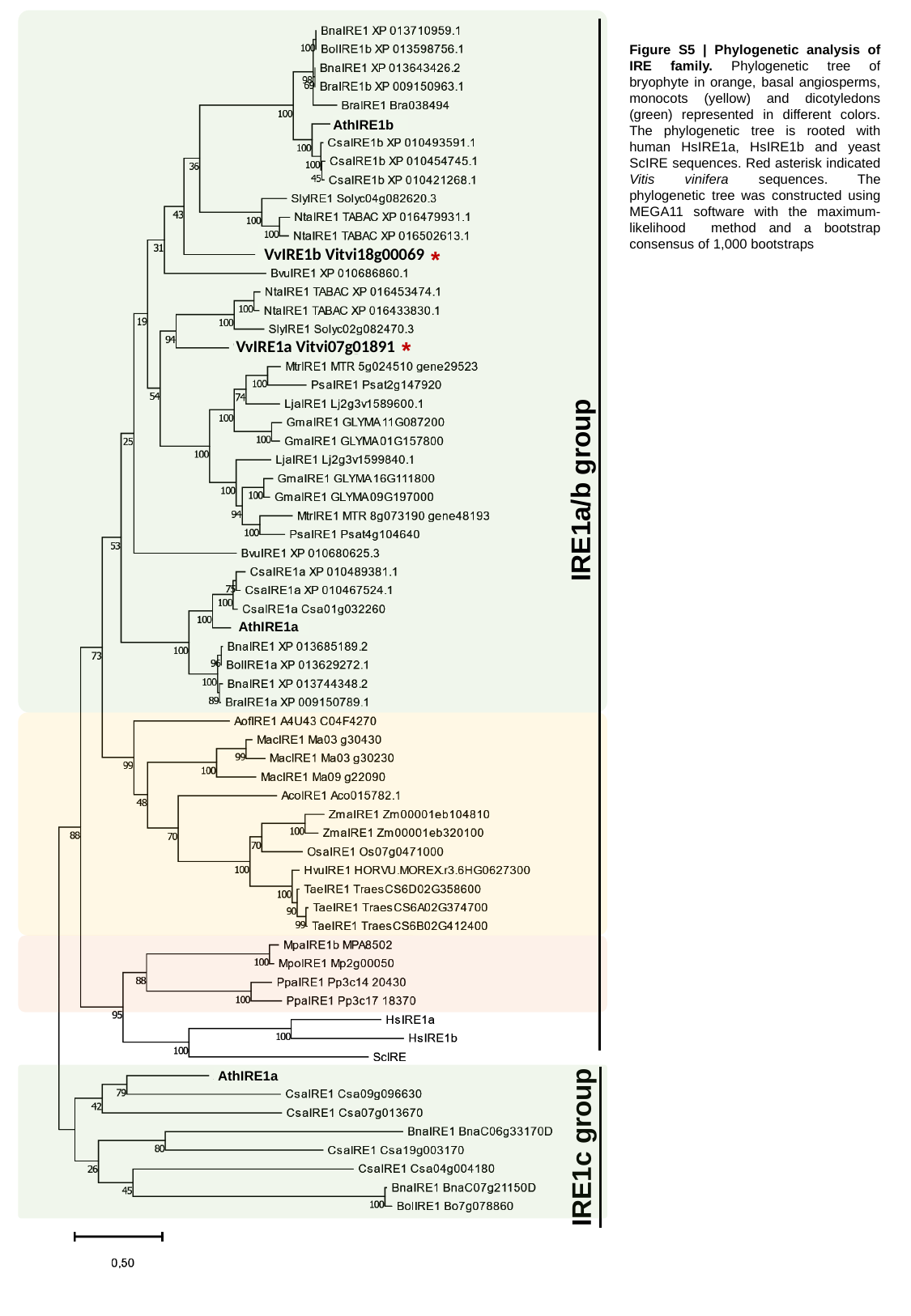

AthIRE1b
*
*
IRE1a/b group
AthIRE1a
IRE1c group
Figure S5 | Phylogenetic analysis of IRE family. Phylogenetic tree of bryophyte in orange, basal angiosperms, monocots (yellow) and dicotyledons (green) represented in different colors. The phylogenetic tree is rooted with human HsIRE1a, HsIRE1b and yeast ScIRE sequences. Red asterisk indicated Vitis vinifera sequences. The phylogenetic tree was constructed using MEGA11 software with the maximum-likelihood method and a bootstrap consensus of 1,000 bootstraps
VvIRE1b Vitvi18g00069
VvIRE1a Vitvi07g01891
AthIRE1a

### Slide 6
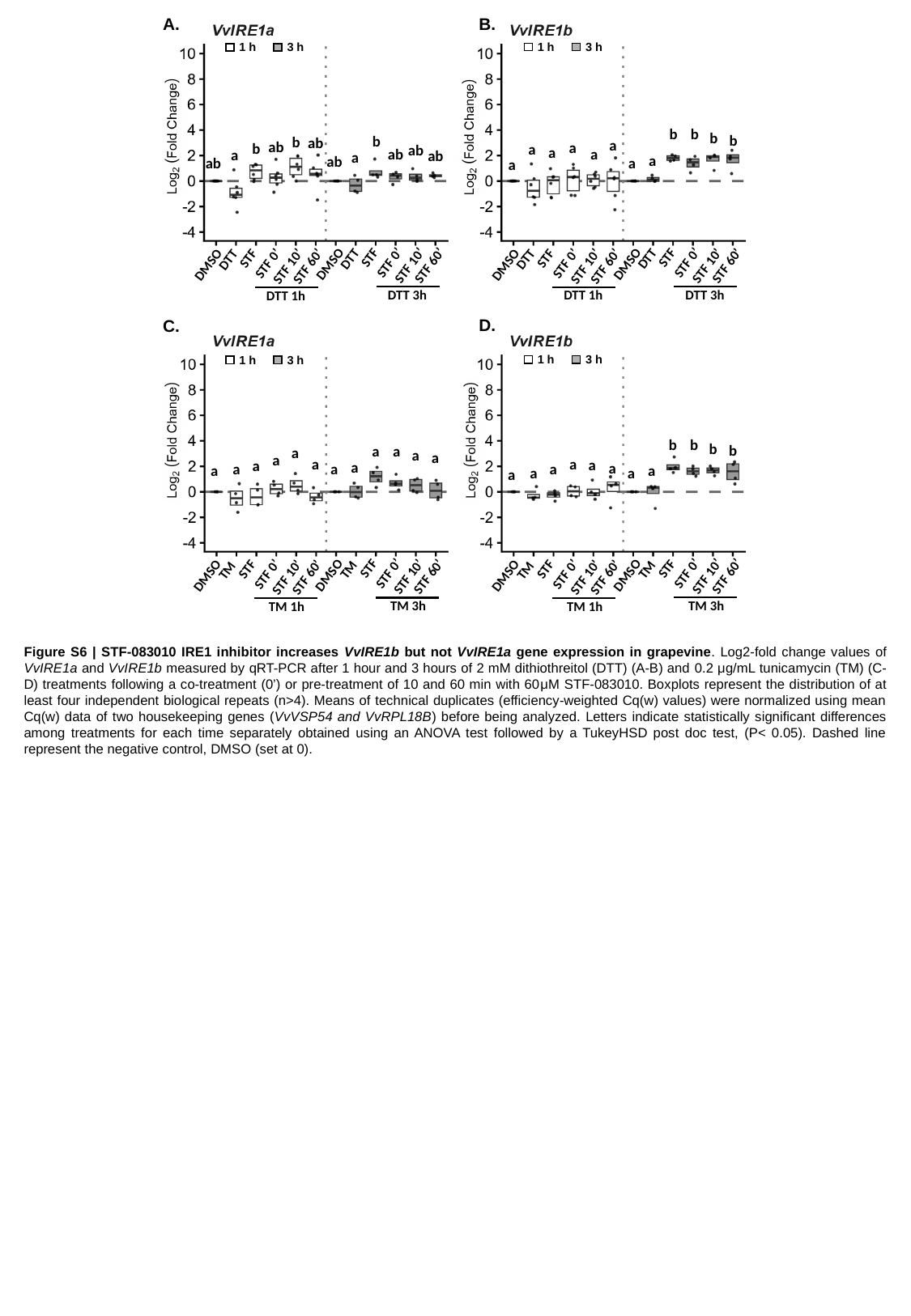

B.
A.
1 h
3 h
1 h
3 h
b
b
b
b
b
b
ab
a
ab
a
b
a
ab
a
ab
a
a
ab
a
a
ab
a
ab
a
STF
DTT
STF 0’
DMSO
STF 10’
STF 60’
DTT 3h
STF
DTT
STF 0’
DMSO
STF 10’
STF 60’
DTT 1h
STF
DTT
STF 0’
DMSO
STF 10’
STF 60’
DTT 3h
STF
DTT
STF 0’
DMSO
STF 10’
STF 60’
DTT 1h
D.
C.
1 h
3 h
1 h
3 h
b
b
b
b
a
a
a
a
a
a
a
a
a
a
a
a
a
a
a
a
a
a
a
a
STF
TM
STF 0’
DMSO
STF 10’
STF 60’
TM 3h
STF
TM
STF 0’
DMSO
STF 10’
STF 60’
TM 1h
STF
TM
STF 0’
DMSO
STF 10’
STF 60’
TM 3h
STF
TM
STF 0’
DMSO
STF 10’
STF 60’
TM 1h
Figure S6 | STF-083010 IRE1 inhibitor increases VvIRE1b but not VvIRE1a gene expression in grapevine. Log2-fold change values of VvIRE1a and VvIRE1b measured by qRT-PCR after 1 hour and 3 hours of 2 mM dithiothreitol (DTT) (A-B) and 0.2 μg/mL tunicamycin (TM) (C-D) treatments following a co-treatment (0’) or pre-treatment of 10 and 60 min with 60μM STF-083010. Boxplots represent the distribution of at least four independent biological repeats (n>4). Means of technical duplicates (efficiency-weighted Cq(w) values) were normalized using mean Cq(w) data of two housekeeping genes (VvVSP54 and VvRPL18B) before being analyzed. Letters indicate statistically significant differences among treatments for each time separately obtained using an ANOVA test followed by a TukeyHSD post doc test, (P< 0.05). Dashed line represent the negative control, DMSO (set at 0).

### Slide 7
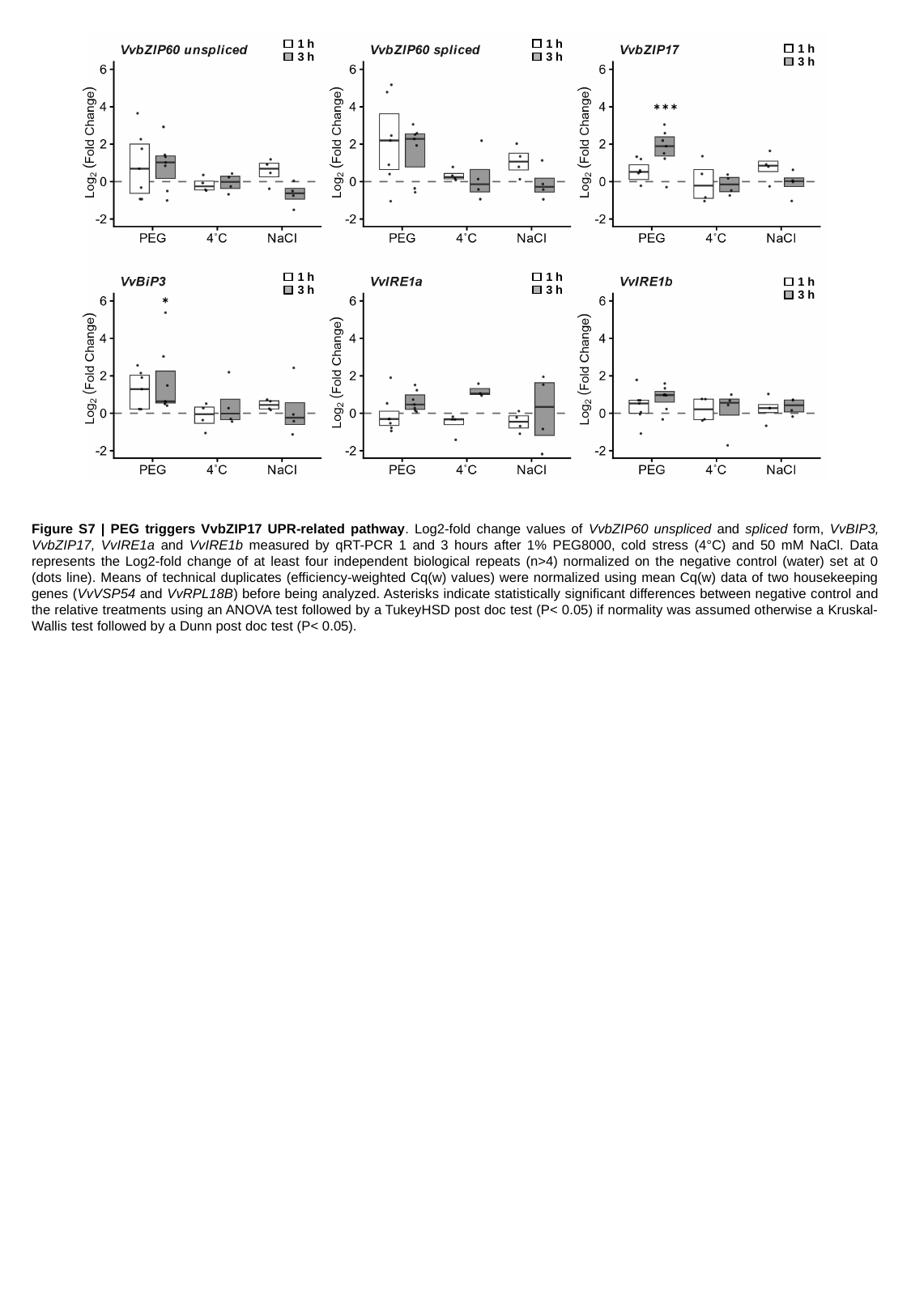

1 h
3 h
1 h
3 h
1 h
3 h
***
1 h
3 h
1 h
3 h
1 h
3 h
*
Figure S7 | PEG triggers VvbZIP17 UPR-related pathway. Log2-fold change values of VvbZIP60 unspliced and spliced form, VvBIP3, VvbZIP17, VvIRE1a and VvIRE1b measured by qRT-PCR 1 and 3 hours after 1% PEG8000, cold stress (4°C) and 50 mM NaCl. Data represents the Log2-fold change of at least four independent biological repeats (n>4) normalized on the negative control (water) set at 0 (dots line). Means of technical duplicates (efficiency-weighted Cq(w) values) were normalized using mean Cq(w) data of two housekeeping genes (VvVSP54 and VvRPL18B) before being analyzed. Asterisks indicate statistically significant differences between negative control and the relative treatments using an ANOVA test followed by a TukeyHSD post doc test (P< 0.05) if normality was assumed otherwise a Kruskal-Wallis test followed by a Dunn post doc test (P< 0.05).

### Slide 8
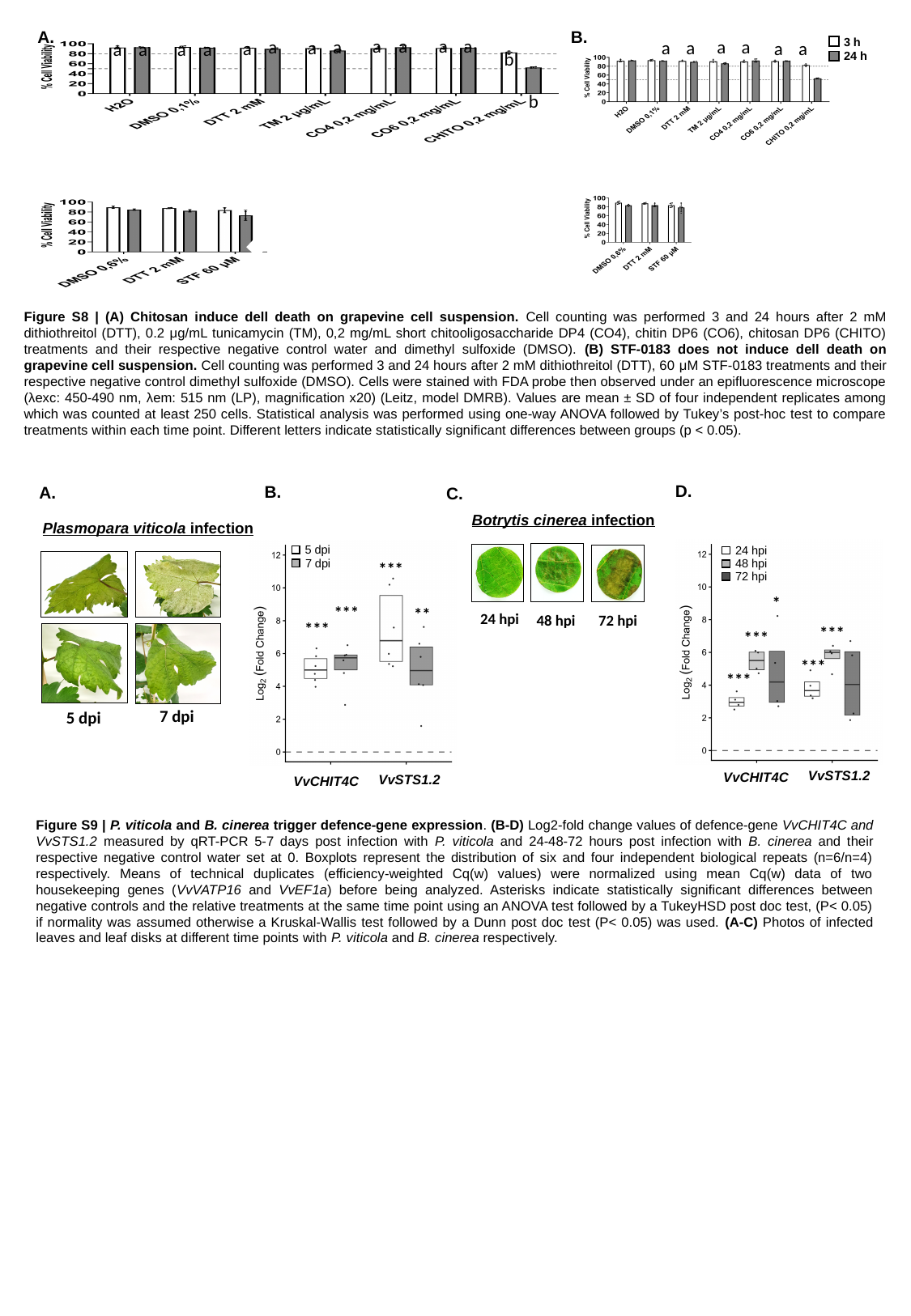

B.
A.
a
a
a
a
a
a
a
a
a
a
a
a
b
b
3 h
24 h
a
a
a
a
a
a
Figure S8 | (A) Chitosan induce dell death on grapevine cell suspension. Cell counting was performed 3 and 24 hours after 2 mM dithiothreitol (DTT), 0.2 μg/mL tunicamycin (TM), 0,2 mg/mL short chitooligosaccharide DP4 (CO4), chitin DP6 (CO6), chitosan DP6 (CHITO) treatments and their respective negative control water and dimethyl sulfoxide (DMSO). (B) STF-0183 does not induce dell death on grapevine cell suspension. Cell counting was performed 3 and 24 hours after 2 mM dithiothreitol (DTT), 60 μM STF-0183 treatments and their respective negative control dimethyl sulfoxide (DMSO). Cells were stained with FDA probe then observed under an epifluorescence microscope (λexc: 450-490 nm, λem: 515 nm (LP), magnification x20) (Leitz, model DMRB). Values are mean ± SD of four independent replicates among which was counted at least 250 cells. Statistical analysis was performed using one-way ANOVA followed by Tukey’s post-hoc test to compare treatments within each time point. Different letters indicate statistically significant differences between groups (p < 0.05).
D.
B.
A.
C.
Botrytis cinerea infection
Plasmopara viticola infection
5 dpi
24 hpi
7 dpi
48 hpi
***
72 hpi
*
***
**
24 hpi
48 hpi
72 hpi
***
***
***
***
***
7 dpi
5 dpi
VvSTS1.2
VvCHIT4C
VvSTS1.2
VvCHIT4C
Figure S9 | P. viticola and B. cinerea trigger defence-gene expression. (B-D) Log2-fold change values of defence-gene VvCHIT4C and VvSTS1.2 measured by qRT-PCR 5-7 days post infection with P. viticola and 24-48-72 hours post infection with B. cinerea and their respective negative control water set at 0. Boxplots represent the distribution of six and four independent biological repeats (n=6/n=4) respectively. Means of technical duplicates (efficiency-weighted Cq(w) values) were normalized using mean Cq(w) data of two housekeeping genes (VvVATP16 and VvEF1a) before being analyzed. Asterisks indicate statistically significant differences between negative controls and the relative treatments at the same time point using an ANOVA test followed by a TukeyHSD post doc test, (P< 0.05) if normality was assumed otherwise a Kruskal-Wallis test followed by a Dunn post doc test (P< 0.05) was used. (A-C) Photos of infected leaves and leaf disks at different time points with P. viticola and B. cinerea respectively.

### Slide 9
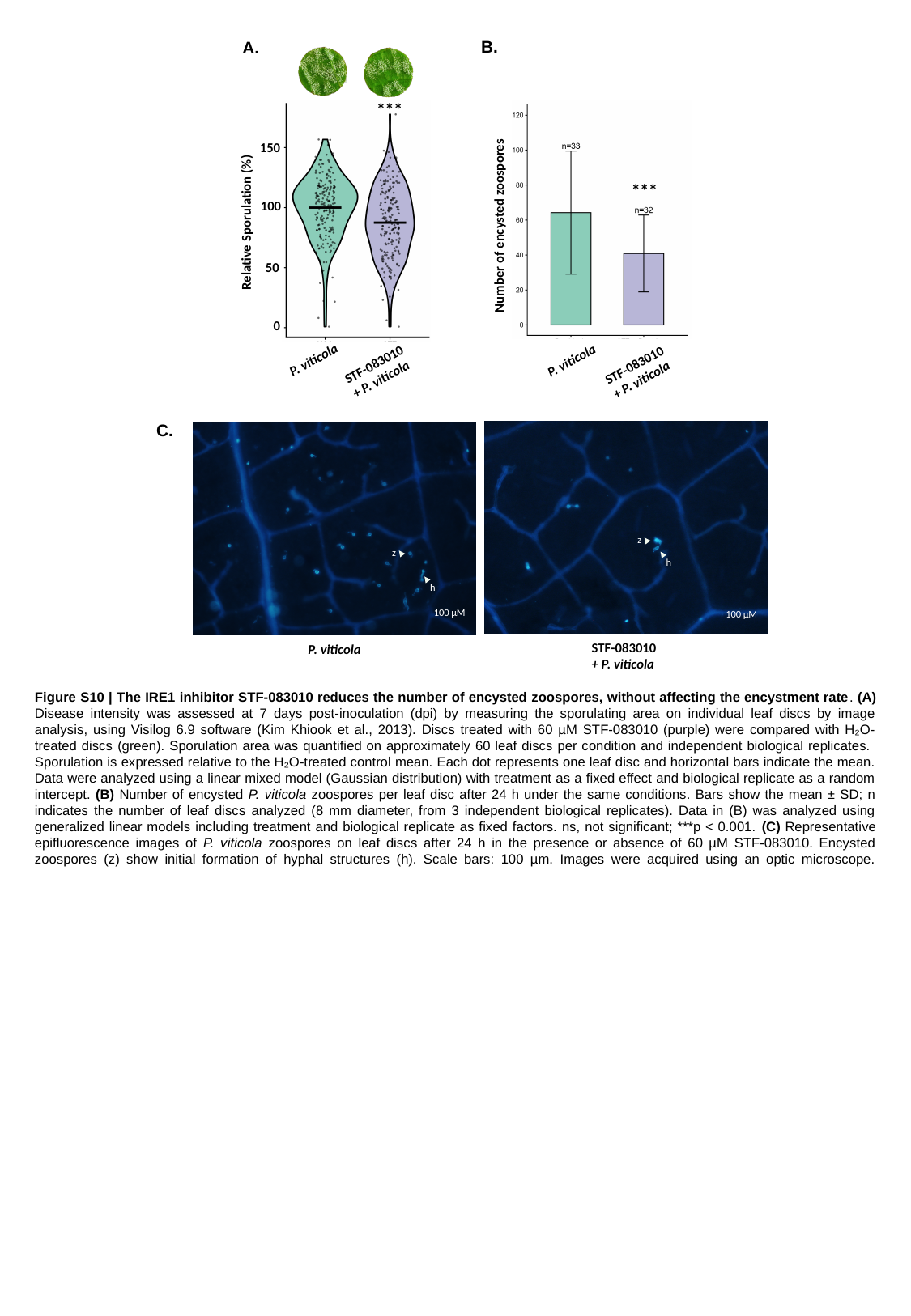

B.
A.
***
150
***
100
Relative Sporulation (%)
Number of encysted zoospores
50
0
P. viticola
P. viticola
STF-083010 + P. viticola
STF-083010 + P. viticola
C.
z
z
h
h
100 µM
100 µM
STF-083010 + P. viticola
P. viticola
Figure S10 | The IRE1 inhibitor STF-083010 reduces the number of encysted zoospores, without affecting the encystment rate. (A) Disease intensity was assessed at 7 days post-inoculation (dpi) by measuring the sporulating area on individual leaf discs by image analysis, using Visilog 6.9 software (Kim Khiook et al., 2013). Discs treated with 60 µM STF-083010 (purple) were compared with H₂O-treated discs (green). Sporulation area was quantified on approximately 60 leaf discs per condition and independent biological replicates. Sporulation is expressed relative to the H₂O-treated control mean. Each dot represents one leaf disc and horizontal bars indicate the mean. Data were analyzed using a linear mixed model (Gaussian distribution) with treatment as a fixed effect and biological replicate as a random intercept. (B) Number of encysted P. viticola zoospores per leaf disc after 24 h under the same conditions. Bars show the mean ± SD; n indicates the number of leaf discs analyzed (8 mm diameter, from 3 independent biological replicates). Data in (B) was analyzed using generalized linear models including treatment and biological replicate as fixed factors. ns, not significant; ***p < 0.001. (C) Representative epifluorescence images of P. viticola zoospores on leaf discs after 24 h in the presence or absence of 60 µM STF-083010. Encysted zoospores (z) show initial formation of hyphal structures (h). Scale bars: 100 µm. Images were acquired using an optic microscope.

### Slide 10
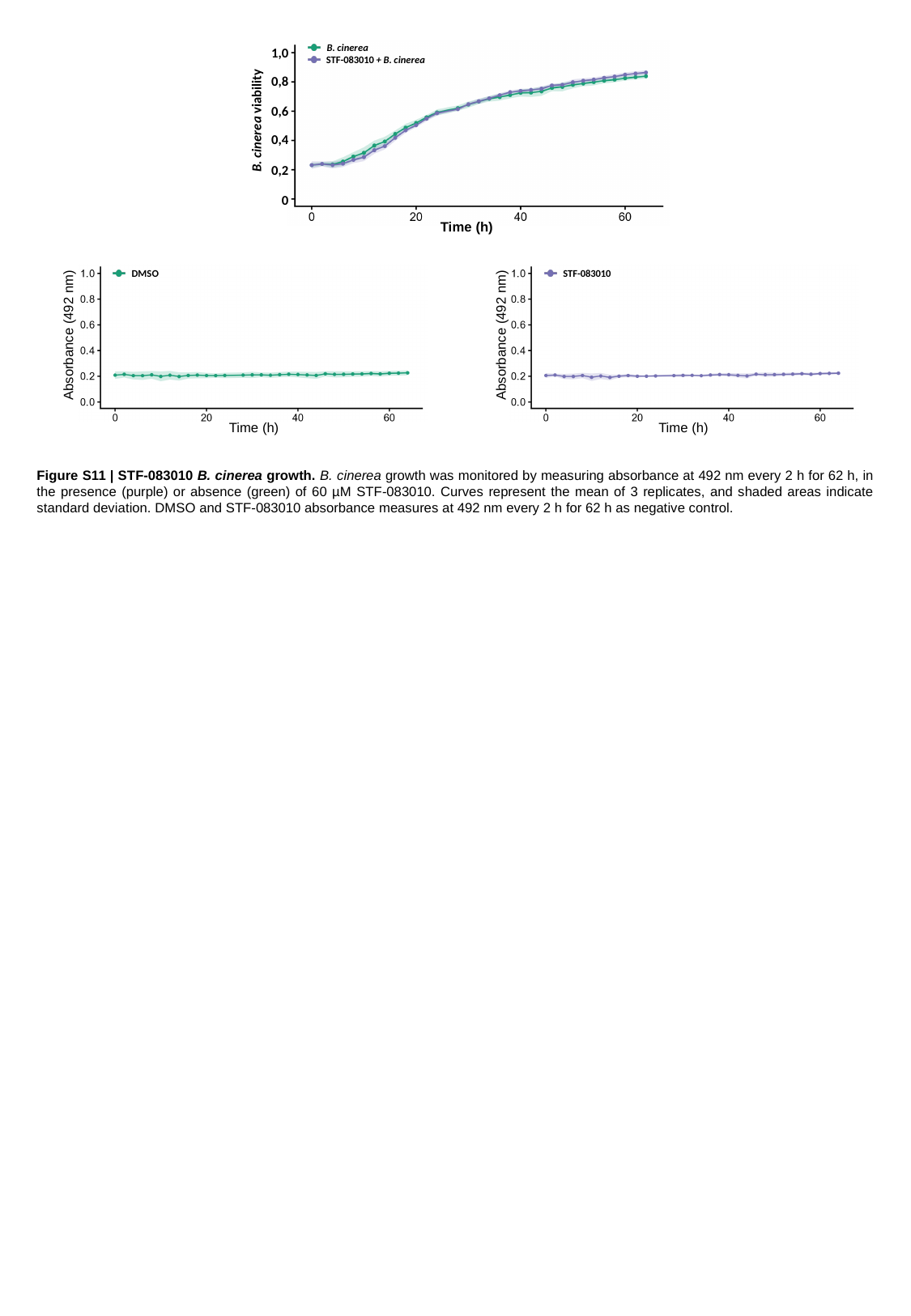

B. cinerea
STF-083010 + B. cinerea
1,0
0,8
0,6
0,4
0,2
0
Time (h)
B. cinerea viability
DMSO
STF-083010
Absorbance (492 nm)
Absorbance (492 nm)
Time (h)
Time (h)
Figure S11 | STF-083010 B. cinerea growth. B. cinerea growth was monitored by measuring absorbance at 492 nm every 2 h for 62 h, in the presence (purple) or absence (green) of 60 µM STF-083010. Curves represent the mean of 3 replicates, and shaded areas indicate standard deviation. DMSO and STF-083010 absorbance measures at 492 nm every 2 h for 62 h as negative control.
